# Renewal theory provides a universal quantitative framework to characterise the continuous regeneration of rotational events in cardiac fibrillation

**DOI:** 10.1101/599142

**Authors:** Dhani Dharmaprani, Madeline Schopp, Pawel Kuklik, Darius Chapman, Anandaroop Lahiri, Lukah Dykes, Feng Xiong, Martin Aguilar, Benjamin Strauss, Lewis Mitchell, Kenneth Pope, Christian Meyer, Stephan Willems, Fadi G. Akar, Stanley Nattel, Andrew D McGavigan, Anand N. Ganesan

## Abstract

**Background:** Cardiac fibrillation is thought to be maintained by rotational activity, with pivoting regions called phase singularities (PSs). Despite a century of research, no clear quantitative framework exists to model the fundamental processes responsible for the continuous formation and destruction of rotors in fibrillation.

**Objective:** We conducted a multi-modality, multi-species study of AF/VF under the hypothesis that PS formation/destruction in fibrillation can be modelled as self-regenerating renewal processes, producing exponential distributions of inter-event times governed by constant rate-parameters defined by the prevailing properties of the system.

**Methods:** PS formation/destruction was studied and cross-validated in 5 models, using basket recordings and optical mapping from: i) human persistent AF (n = 20), ii) tachypaced sheep AF (n = 5), iii) rat AF (n = 4), iv) rat VF (n = 11) and v) computer simulated AF (SIM). Hilbert phase maps were constructed. PS lifetime data were fitted by exponential probability distribution functions (PDFs) computed using maximum entropy theory, and the rate parameter (*λ*) determined. A systematic review was conducted to cross-validate with source data from literature.

**Results:** PS destruction/formation distributions showed good fits to an exponential in all systems (*R*^2^ ≥ 0.90). In humans, *λ* = 4.6%/ms (95%CI,4.3,4.9)), sheep 4.4%/ms (95%CI,4.1,4.7)), rat AF 38%/ms (95%CI,22,55), rat VF 46%/ms (95%CI,31.2,60.2) and SIM 5.4%/ms (95%CI,4.1,6.7). All PS distributions identified through systematic review were exponential with λ comparable to experimental data.

**Conclusion:** These results provide a universal quantitative framework to explain rotor formation and destruction in AF/VF, and a platform for therapeutic advances in cardiac fibrillation.

## INTRODUCTION

Cardiac fibrillation is characterised by aperiodic turbulence of wave propagation(1–3). This can occur in the atria, known as atrial fibrillation (AF), or the ventricles, constituting ventricular fibrillation (VF). The mechanisms by which these arrhythmias are maintained remain incompletely understood, but are of great clinical significance as AF is the most common cardiac arrhythmia in humans, and VF the leading cause of sudden death(4,5).

Although the atria and ventricles possess vastly different geometries, ionic mechanisms and global structures, they share similar electrical wave propagation dynamics(6). A defining characteristic of both AF and VF is the presence of unstable re-entrant circuits that stochastically disappear and spontaneously regenerate during ongoing fibrillation(5,7–18). A key feature of such re-entrant circuits is the presence of a phase singularity (PS) at their pivoting region(19). Although an enormous amount of scientific and clinical effort has gone into the development of strategies to map rotational circuits during fibrillation(16,17,20), a universal quantitative framework to model the processes underlying their formation and destruction is lacking(21).

We reasoned, given the stochastic nature of electrical wave propagation in fibrillation, that the destruction and regeneration of re-entrant circuits may be modelled using renewal theory. Renewal processes are a class of stochastic processes for which the timing of intervals between events can be approximated as statistically independent, identically distributed random variables(22).

We hypothesised that if the rate of PS formation is constant over time, then the time intervals between phase singularity formation and destruction events should follow exponential distributions, implying that these occur via memoryless Poisson point processes with constant rate-parameters. To gain insight into the processes by which AF and VF are sustained, we investigate the statistical distributions of waiting-times for PS formation and destruction with a multi-modality, multi-species study using basket and optical mapping in human, ovine, rat and computer simulated atrial and ventricular fibrillation.

**Figure 1:**
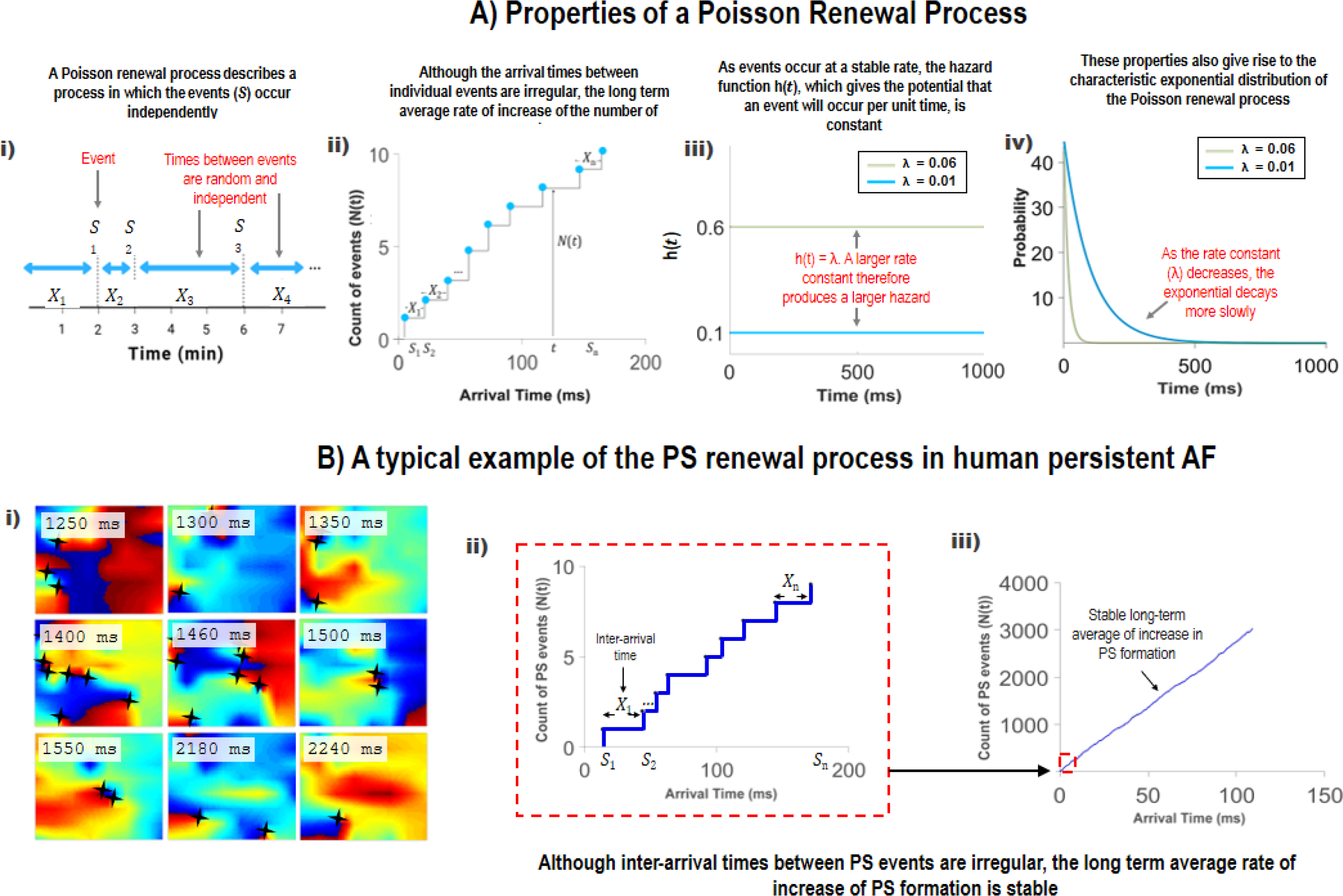
Properties of a Poisson Renewal Process and A typical example of the PS renewal process in human persistent AF. *Figure 1A* A Poisson renewal process is one in which the times between events are independent, but events occur at a long-term constant rate (**i** and **ii**). As events occur at a constant rate, this gives rise to a constant hazard *h*(*t*) (**iii**). The distribution takes the characteristic form of an exponential (**iv**). If data possesses these properties, it implies the generating process is a Poisson renewal process. Examples of Poisson processes in other scientific disciplines include the timing of radioactive decay events, in simple models of survival analysis, or the gating of voltage-gated ion channels. *Figure 1B* **i**) Depicts phase movie snapshots of human persistent AF. In the snapshots, the appearance and disappearance of several PS is shown. The timing of PS is tracked. It can be seen that that PS (marked with *) appear and disappear, although the time intervals between events are irregular. In **ii**), we show a staircase diagram to plot the inter-interval time between new PS formation events. It shows that although the times between new PS events are irregular, the long-term average rate of increase of PS formation (shown by the slope) is stable (**iii**).

## METHODS

A multi-modality, multi species study was performed to analyse basket and optical mapping in: i) human persistent AF, ii) tachypaced sheep AF, iii) rat AF, iv) computer simulated AF and v) rat VF. A systematic review was conducted to cross validate with source data from the literature.

### Human Study Design

The human AF study was a multicentre observational design analysing electrograms acquired prior to ablation. The inclusion criterion was persistent AF undergoing clinically indicated ablation. Patient participation was by informed consent, with recruitment from two centres (Flinders University, Australia), and Hamburg University (Germany). Patient characteristics are in Table 1 and 2, Supplement S1.

### Data Acquisition

#### Human AF

Basket catheter recordings were performed as previously described(23). 64-electrode basket catheters (Constellation, Boston Scientific, MA, 48mm(4mm spacing), 60mm(5mm spacing)) were utilized, based on computed tomographic scan. Unipolar electrogram recordings (1-500 Hz, 2000Hz sampling frequency) were obtained in spontaneous or induced AF lasting ≥5 minutes in the left and right atria. Catheter stability was verified fluoroscopically, and in Velocity (Abbott, IL, USA).

#### Sheep AF

Ovine persistent AF was induced via atrial tachypacing for 16 weeks at ≥ 300 bpm per minute as described(23). Unipolar electrograms were obtained during electrophysiology study, using 64-electrode Constellation catheters (48mm). Electrograms were filtered from 1 Hz to 500 Hz, sampled at 2 kHz on NavX (Abbott, IL, USA). ≥5 minutes LA and RA recordings were obtained.

#### Rat AF

Optical mapping in a rat AF model was performed as previously described (S12)(19,24,25). The heart was excised and perfused in Langendorff mode with Krebs solution at 30 ml/min and 37°C. After 30 min for stabilization and electrical/mechanical decoupling with blebbistatin (15 μM), the heart was loaded with di-4-ANEPPS (Biotium, Inc., Hayward, California). A charge-coupled device (CardioCCD, RedShirtImaging, LLC, Decatur, Georgia) was used to record RA free wall fluorescence at 1 kHz. Bipolar electrodes were used to pace the right atrial appendage. Optical maps were obtained during AF induced by 1.5 × threshold current 2-ms stimulation at a reducing cycle length from 150ms to 20 ms.

#### Rat VF

Optical mapping in a rat VF model was performed as previously described(26). Rat hearts were excised and transferred to a Langendorff apparatus, and retrogradedly perfused through the aorta with oxygenated Tyrodes solution. Perfusion pressure was maintained at ~60 mm Hg by adjusting perfusion flow. Hearts were regulated at 37° C. Volume conducted electrocardiograms (EGMs) were recorded. Background fluorescence intensity was measured periodically in 1 minute intervals, using a 6400 pixel charge coupled device.

#### Computer Simulated AF

Computer simulations were based on the Ten Tusscher-Panfilov model on 2-D, isotropic square grids (80×80)(23). Model calculations are in S2. Three AF-type scenarios were generated: i) stable spirals surrounded by moderately disorganised wavebreak, ii) stable spirals surrounded by highly disorganised wavebreak, iii) multiple wavelet re-entry without stable spirals and iv) stable spiral. These scenarios were selected to understand the effect of spiral breakup on PS destruction and formation.

### Cleaning, Filtering and Sinusoidal Recomposition

Signal processing was performed similarly to previous studies(27). Unipolar electrograms, surface ECG, and 3D data were exported from NavX. QRS subtraction was performed (S3)(28). Signals were filtered using a 4th order Butterworth fitted with a 1 to 30 Hz band pass filter applied in forward and reverse mode(27). A Hanning window was applied for edge tapering, and the Welch power spectral method applied. The Fourier dominant frequency was used as the wavelet period for sinusoidal wavelet recomposition(29).

#### Hilbert Phase Mapping and Phase Singularity Detection

Instantaneous phase for each electrogram was computed by applying the Hilbert transform (S3)(27,29).APS appears when the phase around a given point progresses through from to -π to +π(30). PS detection was performed using the double-ring method to avoid noise-related issues (S3)(31). For sensitivity analysis, the Iyer and Gray line integral method was also performed(32).

### PS look-up table

To determine PS lifetime and PS inter-formation event times, we created a look-up table indexing onset time, offset time and electrode location for each new PS detection. A new PS detection event was defined as the detection of a PS at an electrode and its surrounding 1-electrode neighbourhood for ≥3 consecutive frames. The look-up table enabled computation of histograms for: (i) PS lifetimes and (ii) inter-formation event times.

### PS lifetime distribution as an index of PS destruction

For each epoch, observed experimental PS lifetime data was fitted with an exponential distribution using non-linear least-squares regression (Figure 3A,3F), and the PS destruction rate (λ) estimated. The adequacy of fit was determined using the R-squared value (*R*^*2*^). To validate, PS data was fitted using maximum likelihood and adequacy of fit evaluated using the chi-squared (χ^2^) goodness of fit test (S4).

### Cumulative distribution of inter-formation times as an index of new PS formation

To evaluate the contribution of stochastic processes to PS formation, we calculated *R*^*2*^ of the cumulative distribution functions (CDF). The exponential CDF can be given by:

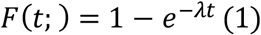

where λ is the rate parameter, and *t* the time between formation events. Maximum likelihood was used for cross-comparison (S4).

### Sensitivity analyses

To cross-validate, sensitivity analyses were conducted. First, electrode permutation was performed to destroy spatial correlations and assess the sensitivity of PS detection methods to noise. For each epoch, an electrode permutation series was created by randomly permuting electrodes. PS detection and computation of λ was performed on each permuted dataset. λ values were compared to those derived from non-permuted observed experimental data (S5).

A second PS detection method, the line integral approach by Iyer and Gray(32), was also used to verify of the specificity of phase singularity detection. λ values were determined and compared (S6).

### Comparison with MaxEnt Predicted Distribution

MaxEnt is a rigorously proven principle of statistical inference allowing prediction of the least-biased (most purely random) underlying probability distribution, given prior constraints. MaxEnt has been applied in physics(33), neuroscience(34), and ecology(35). For a waiting-time process, with sample mean as a constraint, MaxEnt predicts an exponential probability distribution function (PDF):

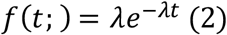

where λ is the rate. For the PS-formation function,*t* represents time between formation events. For PS-destruction, *t* represents PS lifetime. A step-by-step guide is in S7.

If the observed data distribution matches that predicted by MaxEnt, it implies the generating process can be accurately modelled as purely stochastic, except for the constraint of the mean rate. We compare the distributions of observed waiting-times for PS formation and destruction to those predicted by MaxEnt to validate whether these processes are truly stochastic and maximally random.

### Comparison with PS Data in Other Studies

For further cross-validation with published source data, we conducted a systematic review of PubMed English language medical literature to study the distribution of PS lifetimes. Methods and search and inclusion criteria are outlined in S8.

## RESULTS

### PS Destruction in Human and Animal Cardiac Fibrillation

PS lifetime distributions for all human persistent AF cases showed an exponential distribution (Figure 2), with R-squared values (*R*^*2*^) ≥ 0.90. A summary of data fitting results is shown in Table 1 (Supplement S4). PS lifetime hazard functions *h*(*t*) demonstrated a constant λ rate for all cases (Figure 2B). The mean PS destruction rate (λ) was 4.6%/ms±1.5 (95%CI, 4.3, 4.9), translating to a PS half-life (t_1/2_) of 15 ms (95%CI, 14, 16). There were no statistically significant differences (P=0.18) in PS destruction λ between the LA (4.5%/ms (95%CI, 4.1, 4.8)) and RA (4.9%/ms (95%CI, 4.3, 5.4)(Supplement S9).

**Figure 2:**
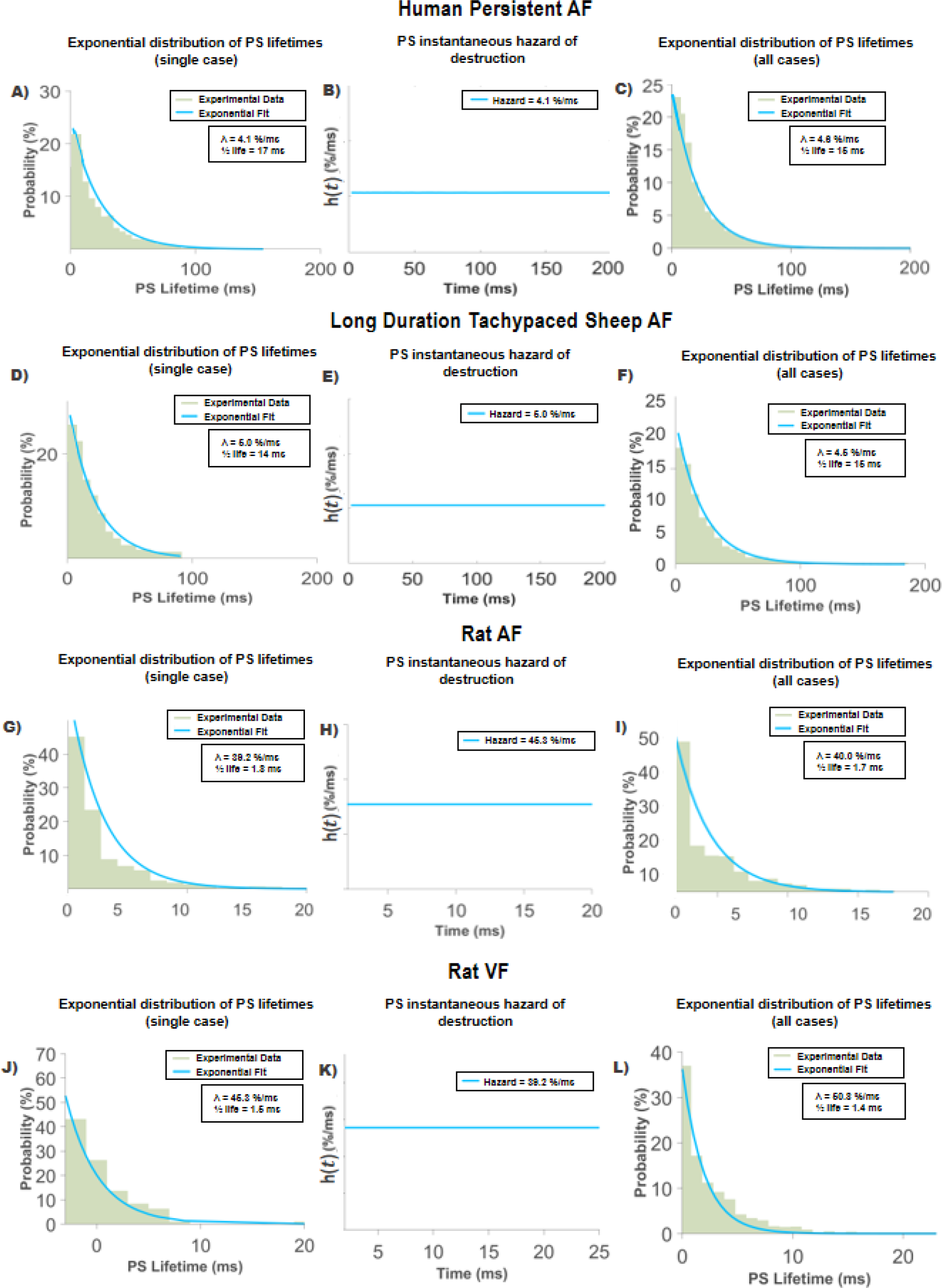
PS destruction occurs via a Poisson renewal process. PS lifetime distributions in all model systems were exponential. **Persistent Human AF (2A-C):** PS lifetime distributions for **2A**- single case; **2C**- all patients (n=20) plotted together. **2B** instantaneous hazard shows constant rate. **Sheep tachypacing persistent AF (2D-F):** PS lifetime distributions for **2D**- single case; **2F**- all tachypaced sheep (n=18). **Rat AF (3G-I):** PS lifetime distributions for **2G**- single case; **2I**- all rats (n=4). **Rat VF (3J-L):** PS lifetime distributions for **2J**- single case; **2L**- all rats (n=11). Together, these properties are consistent with a stochastic Poisson renewal process.

PS destruction in sheep tachypaced sustained AF was similar to humans, with PS lifetime distributions showing good fits to the exponential distribution (*R*^*2*^ ≥ 0.90 (Table 2, S4). In sheep, the mean λ was 4.4%/ms±1.4 (95%CI, 4.1, 4.7), corresponding to a PS half-life (t_1/2_) of 16 ms (95%CI, 15, 18). Mean λ in sheep did not differ significantly between LA (4.5%/ms (95%CI, 4.02, 5.1)) and RA (4.22%/ms (95%CI, 3.8, 4.6)) (P=0.20) (S9).

Rat AF data demonstrated PS lifetime distributions that were well fitted by an exponential distribution (*R*^*2*^ ≥ 0.90)(Table 3, S4), with a mean λ of 38%/ms±6.6 (95%CI, 22, 55). (t_1/2_) was 1.8 ms (95%CI, 1.2, 3.1). Faster λ may be accounted for by the differences in action potential dynamics of the rat in comparison to larger mammals.

In rat ventricular fibrillation, PS lifetime curves also showed good fits to an exponential distribution (*R*^*2*^ ≥ 0.92)(Table 4, S4). Similarly to rat AF, faster λ was observed (λ = 46%/ms ±21 (95%CI, 31, 60), (t_1/2_) was 1.5 ms±3.3 (95%CI, 2.2, 5.7)).

### PS Formation in Human and Animal Cardiac Fibrillation

Similarly to PS destruction, PS formation probability density functions (3A) showed exponential distributions (*R*^*2*^ ≥ 0.90 (Table 5, S4)). For interpretability, we also provide the cumulative density function to show the probability that a new PS has formed after an interformation event time of *t* (Figure 3, Equation 1). Mean λ was 4.6%/ms±1.1 (95%CI, 4.3, 5.0), translating to a t_1/2_ of 15 ms (95%CI, 14, 16). λ in the LA (4.9%/ms (95%CI, 4.5, 5.4)) was not different to the RA (4.1%/ms (95%CI, 3.6, 4.6) (P=0.28) (S9)).

**Figure 3:**
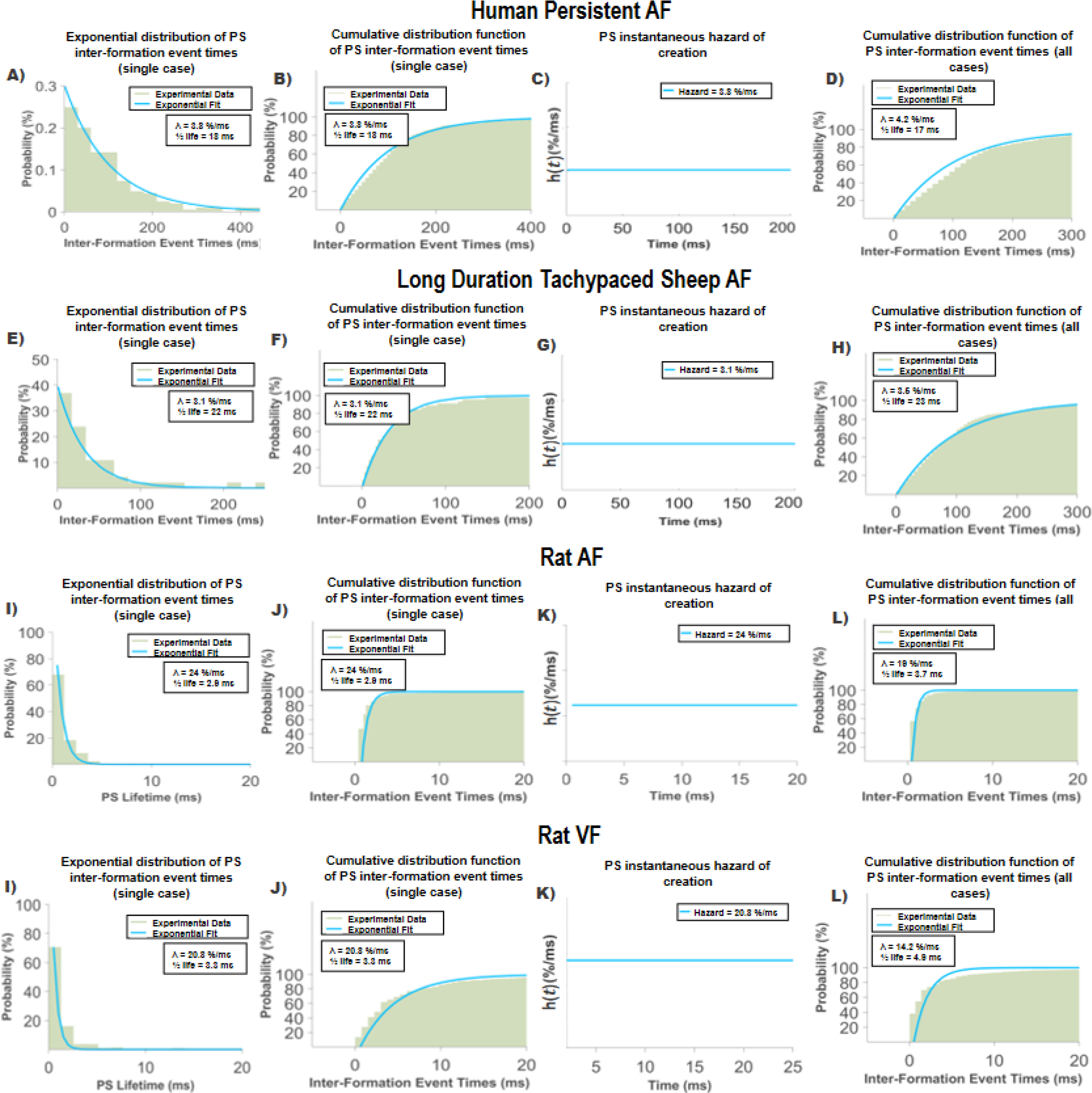
PS formation occurs via a Poisson renewal process. PS inter-formation event time distributions were exponential in all models. **Human persistent AF (3A-D)** PS inter-formation event time distributions for **3A,B**- single case; **3D**- all patients (n=20). **3C** instantaneous hazard shows constant rate. **Sheep tachypacing persistent AF (3E-H)** PS inter-formation event time distributions for **3E,F**- single case; **3H**- all tachypaced sheep (n=18). **Rat AF (3I-L):** PS inter-formation event time distributions for **3E,F**- single case; **3H**- all rats (n=4). **Rat VF (3I-L):** PS inter-formation event time distributions for **3I,J**- single case; **3L**- all rats (n=11).

Tachypaced sheep curves also showed good fits with exponential distributions in all cases (*R*^*2*^ ≥ 0.90 (Table 6 S4). Mean λ was 4.6%/ms±1.4 (95%CI, 4.3, 4.8) and t_1/2_ equal to 15 ms (95%CI, 14, 16). λ for PS formation in the LA (LA: 4.5%ms/s (95%CI, 5.1, 6.7) was not different to the RA (4.6%ms/s (95%CI, 5.3, 6.9))(P = 0.79).

Rat AF also showed good fits with an exponential distribution for all cases (*R*^*2*^ ≥ 0.90). Mean λ was 33%/ms ± 8.8 (95%CI, 11, 55) and t_1/2_ equal to 2.1 ms (95%CI, 12, 6.3). Again, λ was faster in comparison to human and sheep (Table 7, S4).

In rat VF, mean λ was 38%/ms±24 (95%CI, 22, 55) and t_1/2_ equal to 1.8 ms (95%CI, 1.3, 3.2). All cases showed good fits with an exponential distribution (*R*^*2*^ ≥ 0.91) (Table 8, S4).

### PS Formation and Destruction in Computer Simulated AF

Figure 4 shows the distribution of PS lifetimes in the Ten-Tusscher model of AF in 3 different scenarios:i) stable spirals surrounded by highly disorganised wave breakup, ii) stable spirals surrounded by moderately disorganised breakup and iii) wavelet re-entry without stable spirals. All SIM AF PS lifetime distributions showed good fits to an exponential distribution, irrespective of the presence of stable spirals (highly disorganised: *R*^*2*^=0.97; moderately disorganised: *R*^*2*^=0.99; wavelet re-entry: *R*^*2*^=0.99;).

**Figure 4:**
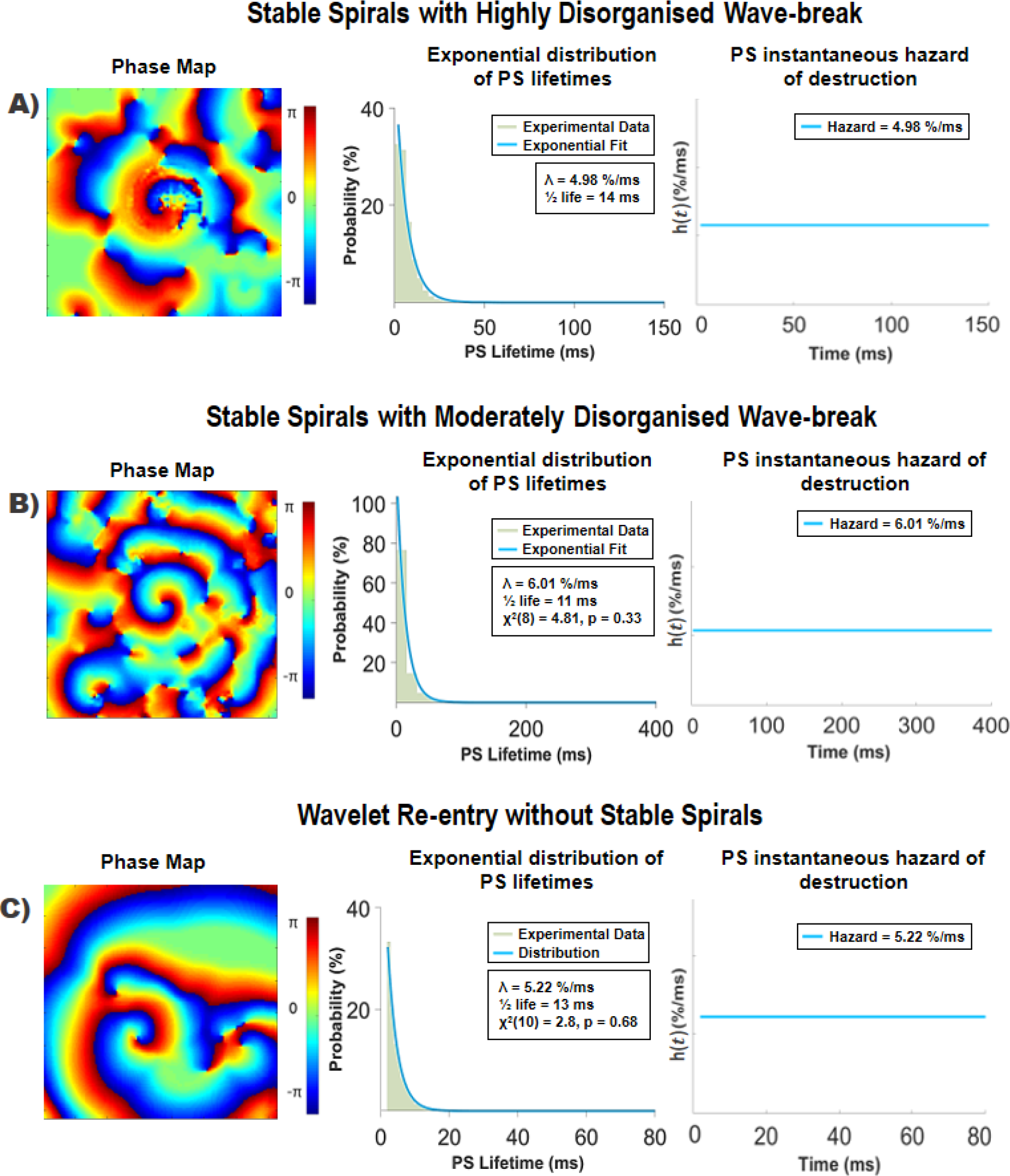
PS lifetime distributions under different scenarios of AF. PS lifetime distributions were exponential, irrespective of the presence of a stable rotors. Hazard rates were constant. Simulations depicting spirals with moderately disorganised wave break yielded largest λ and longer lasting PS.

### Comparison with Line Integral PS Detection Method

For sensitivity analysis, datasets were subjected to repeat analyses using the line integral PS detection method(32). PS lifetime and inter-formation event time curves showed exponential distributions (S6), consistent with findings using the double-ring method. In humans λ was equal to 4.5%/ms (95%CI, 3.8, 4.3; t_1/2_: 15ms (95%CI, 16, 18)). In sheep AF, λ was 3.7%/ms (95%CI, 3.4, 4.0; t_1/2_: 19ms (95%CI, 17, 20)). PS formation was similar, with λ in humans equal to 4.8%/ms (95%CI, 4.6, 5.1) and in sheep, 4.4%/ms (95%CI, 4.2, 4.7).

### Comparison with Maximum Entropy Predicted Distribution

PS lifetime distributions for PS destruction returned similar experimental λ and MaxEnt derived λ in both humans and sheep (Human: 4.9%/ms (95%CI, 4.6, 5.3); Sheep: 4.6%/ms (95%CI, 4.2, 5.0)) (S7), which were strongly correlated (Human R^2^: 0.99; Sheep R^2^: 0.99). PS formation demonstrated similar results (Human: 4.3%/ms (95%CI, 4.0, 4.5); Sheep: 3.7%/ms (95%CI, 3.3).

### Comparison with Experimental Observations from Previously Published Studies

We identified 1782 references, with 23 retrieved for full-text review. We identified 6 articles producing a phase singularity distribution histogram. The results of the search are outlined in S8. PS lifetime histograms from published papers are reproduced in S8, showing that these distributions were consistent with an exponential form (Figure 5, Table 1 S8).

**Figure 5:**
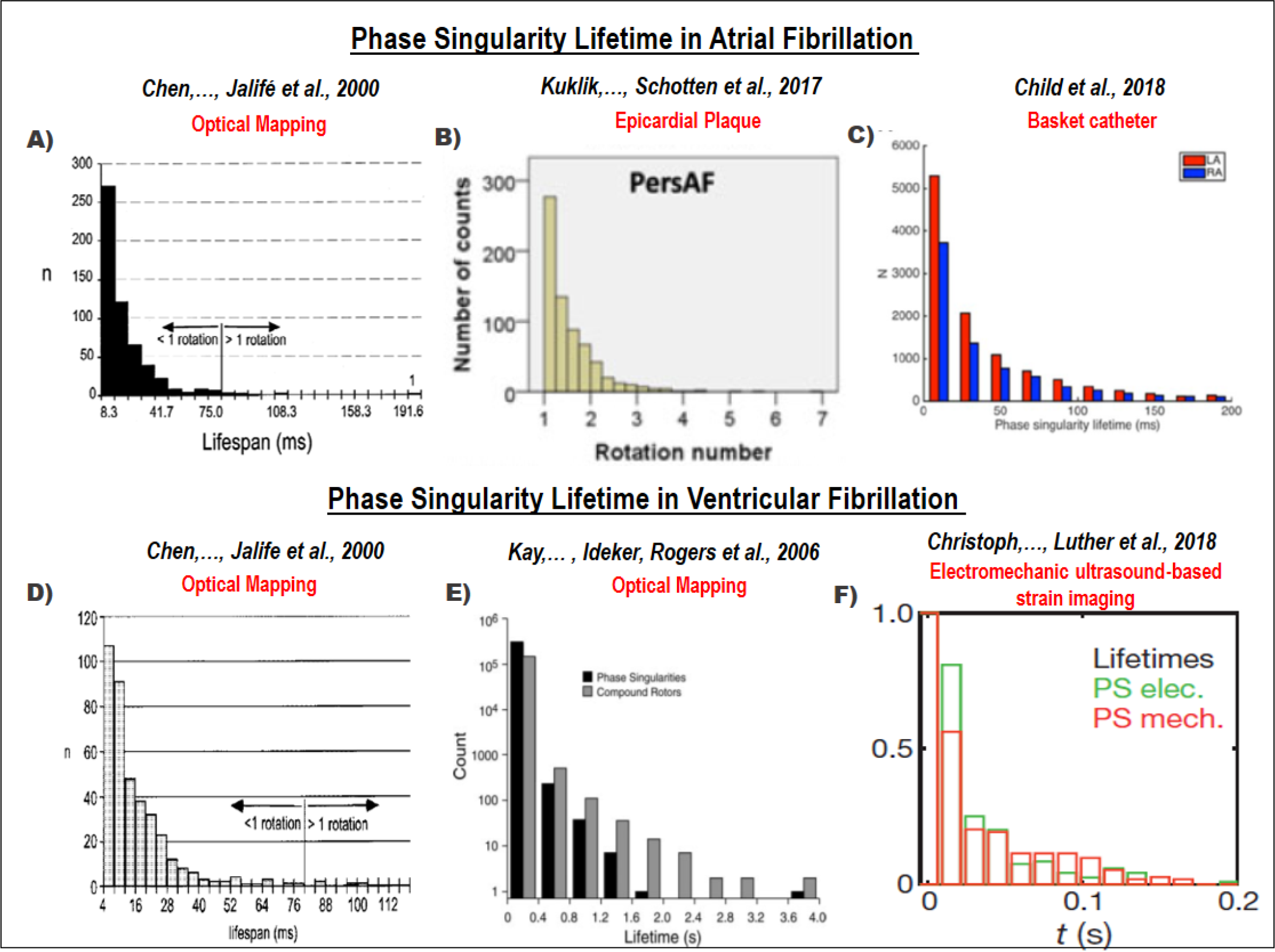
Phase Singularity Lifetime Histograms from Previously Published Studies Consistently Demonstrate Exponential Distributions Irrespective of Mapping Modality. A systematic review was conducted, and 6 articles reporting phase singularity distribution histograms analysed **(5A-F)**. Histograms in both AF and VF consistently demonstrated an exponential distribution of PS lifetimes, irrespective of the mapping approach used (basket catheter, optical mapping, epicardial plaque and electromechanical ultrasound-based mapping).

## DISCUSSION

The continuous appearance and disappearance of unstable re-entrant circuits is a distinguishing feature of cardiac fibrillation. Here, we show that PS lifetimes and inter-formation event times during AF and VF show exponential distributions, consistent with the notion that PS disappear and are regenerated by Poisson point processes. The statistical distribution of PS lifetimes and formation processes are consistent with those predicted by the principle of MaxEnt, suggesting these processes can be accurately modelled as a self-regenerating renewal process. The results provide a universal quantitative framework to characterise the formation and destruction of PS in cardiac fibrillation, and indicate that paradoxically during periods of sustained AF and VF, the system dynamics are conserved, despite the aperiodic turbulence of wave propagation. The rate constants for PS formation and destruction may provide useful parameters to model stochastic elements of fibrillation dynamics.

### Unstable re-entrant circuits - a historical context

The observation of unstable re-entrant circuits in cardiac fibrillation has been a consistent finding throughout the history of the field. Unstable local re-entrant circuits were observed in early works by Garrey(36), who noted the presence of “…a series of ring-like circuits of shifting location and multiple complexity”. Later, computational models of Moe and Abildskov also described a similar phenomenon, describing the “…numerous [unstable] vortices, shifting in position and direction like eddies in a turbulent pool.” Similar results have been in subsequent more physiological computational models. Re-entrant activity has also been a feature of multiple epicardial mapping studies, in both experimental models and in humans(11,37). Similar observations have been observed using optical mapping, where even in cholinergic AF models in which some rotors were more sustained, the majority of PS were considered unstable(13,14,38).

Thus far, the characterization of these processes has predominantly been qualitative. Indeed, the absence of quantifiable predictions based on the multiple wavelet theory has been a criticism levelled at Moe’s original conceptualisation(21). Although multiple reports have observed the stochastic nature of the formation and disappearance of re-entrant circuits, the statistical processes underlying these events have not been well studied.

An interesting feature of the previous literature is that exponential distributions of PS lifetimes are observable in many published data from AF (14,27,31) and VF(14,39,40). However, in these studies the connection of the distribution to the underlying data generating process was not explored. Collectively, however, this provides support for results of the current study.

### Significance of the exponential distribution, and relationship to Poisson point processes

The reason the exponential distribution is so important, is that it gives a window into the kinetic properties of the underlying process(35,41). The exponential distribution occurs in stochastic physical and biological Poisson point processes that are governed by constant rate parameters, such as radioactive decay. The reason this is important in the context of cardiac fibrillation, is that it means PS formation and destruction events occur in a similar stochastic fashion, and can be well-modelled as Poisson point processes. An important notion here is that the Poisson process is *memoryless*. The exponential is the only continuous distribution characteristic of a memoryless process. In simple terms, this means the waiting time for the next event to happen is independent of any history of prior events, and is therefore temporally stable.

### AF and VF as regenerative renewal processes

Poisson point processes are the archetypal example of a renewal process, in which the inter-event waiting times are independent, but the long-term rate is constant. A characteristic of renewal processes is the idea of *regeneration*. This is consistent with a model of AF and VF in which new PSs are in a constant self-regenerating cycle of formation and destruction. Mechanistically, the results may perhaps provide a bridge between the multiple wavelet and rotor theories of fibrillation, by suggesting that continuous generation and destruction of PS (e.g wavebreaks in Moe’s theory), can be conceptualized as a memoryless, stochastic processes, strictly governed by measurable rate parameters.

### Origins of the exponential limiting distribution - applying the principle of maximum entropy

An important question is the origin of the exponential distribution of PS formation and destruction. The phenomenon of microscopic uncorrelated stochastic processes adding to macroscopically stable limiting distributions is a constant motif repeated throughout nature(35,41). This occurs because if the system is averaged over long periods of time, then fluctuations in dynamics of natural systems tend to summate towards recognisable distributions. The principle of MaxEnt(42) has been used in a variety of fields including neuroscience(43) and ecology(43) to enable prediction of the least biased (most stochastic) distribution of a sample. Demonstration that the observed data fits the MaxEnt predicted distribution shows that lifetime processes of PS formation and destruction can be modelled as stochastic processes.

### Implications for overall cardiac fibrillation system dynamics

The switch from sinus rhythm into fibrillation has been considered a form of chaos(44,45). Such systems may have the property of being dissipative, in the sense that the long-term trajectory through the system phase space is unstable, or conservative, so that the long-term behaviour of the system dynamics in phase space is conserved(46). By demonstrating that the processes of PS destruction and formation are regenerative renewal processes, it suggests that this aspect of sustained AF and VF is a conservative, rather than dissipative, nonlinear system.(46) This is consistent with the clinical behaviour of AF, which in persistent forms can last for decades. The nature of how AF becomes more persistent, and how terminations in AF and VF occur, remain questions that will be objectives of future studies.

An interesting result from the computational models analysed was that exponential PS lifetime distributions were observed in the presence of a stable and unstable spiral wave, suggesting these distributions are a measure of the stochastic element of the spiral breakup process.

### Future Directions

By recasting the formation and destruction of rotational events as renewal processes, the results provide a universal theoretical basis for the modelling of rates of PS destruction and formation, which may be related to the persistence and termination of the arrhythmia. These rate constants are easily measurable statistical parameters that allow the fibrillatory process to be characterised.

Future research efforts will be directed to understand how AF and VF terminate in a more systematic way, and to determine the effects of autonomic nervous system regulation, substrate modification, pharmacological therapy and ablation strategies on renewal process parameters in relationship to successful termination or prevention of fibrillation, in order to obtain new mechanistic insights.

## CONCLUSION

We demonstrate that PS destruction and formation in cardiac fibrillation are self-regenerating renewal processes. These results provide a potentially powerful universal quantitative framework to explain rotor formation and destruction in AF and VF, and a platform for therapeutic advances in cardiac fibrillation.

## Acknowledgement of Funding

This work was supported by the National Health and Medical Research Council of Australia Project Grant (1063754) and National Heart Foundation of Australia (101188).

## Abbreviations

AF: Atrial fibrillation
VF: Ventricular Fibrillation
PS: Phase Singularities
EGM: Electrocardiogram
MaxEnt: Maximum Entropy

