## Supplementary Materials for "Renewal theory provides a universal quantitative framework to characterise the continuous regeneration of rotational events in cardiac fibrillation"

#### S1 – Patient Baseline Characteristics

**Table 5: Flinders Medical Centre Cohort Patient Baseline Characteristics**

| | |
|---|---|
| Age | 62±9 |
| Gender | |
| Male | 83%(10) |
| Female | 17%(2) |
| BMI | 28.7±3.8 |
| $CHA_2DS_2VaSC$ | 1.6±1 |
| AF type | |
| Persistent | 80%(8) |
| Paroxysmal | 20%(2) |
| Echocardiographic Parameters | |
| LVEF (%) | 59±7 |
| LA area ($cm^2$) | 25±4 |
| E/E’ | 8.4±1.6 |

**Table 6: University Medical Centre Hamburg Cohort Patient Baseline Characteristics**

| | |
|---|---|
| Age [y] | 60±10 |
| Male [%] | 12/13 |
| BMI [$kg/m^2$] | 29±5 |
| LA diameter [mm] | 48±6 |
| Reduced LV EF (40-55%) | 2/13 |
| Hypertension | 3/14 |
| Diabetes | 0/14 |
| Stroke/TIA | 1/14 |

#### S2 – Mathematical Model of Fibrillatory Activity

One of the mechanisms of wave conduction driving AF is a presence of dynamically unstable area (e.g. high degree of electrical alternans) causing continuous wave break and fibrillatory conduction. Dynamically unstable areas would continuously disrupt wave conduction in the surrounding, generating short-lived pivoting waves. In order to model this scenario, we used a Panfilov-Tusscher model which, due to its simple formulation, allows straightforward control of dynamical stability of the wave conduction.

We used rectangular grid of 100x100 diffusively coupled simulation nodes (diffusion coefficient set to 0.2). Space step was set to 0.6 and time step to 0.02. System was integrated using forward Euler scheme with no-flux boundary conditions. In order to model an inhomogeneous substrate for AF we varied parameter μ2 in the model which strongly influences steepness of the restitution curve. We introduced a random distribution of μ2 which led to the presence of localized areas of high dynamical instability (leading to spiral wave breakup) surrounded by dynamically stable areas.

Based on study by ten Tusscher and Panfilov21, we used following parameters of the model: a=0.1, k=8, ε=0.01, b=0.1, μ1=0.2, μ2=0.2. In order to induce dynamical instability, we varied value of μ1 to 0.02 and 0.2. We used this simplified model only to illustrate the link between dysynchrony and regions maintaining AF due to electrical instability. There are several mechanisms thought to play an important role in AF maintenance (e.g. fibrosis, automaticity, heterogeneity of effective refractory period). However, exploration of all potential factors is beyond the scope of this work and we decided to focus on selected one (heterogeneity of electrical alternans) to demonstrate how dyssynchrony can point to regions critical to AF maintenance. Since we did not aim to model detailed electrophysiology of the atria but rather show specific mechanism, we used arbitrary units.

#### Extracellular EGM Calculation

Bipolar EGMs from a two-dimensional sheet based on the Tusscher-Panfilov phenomelogical model were investigated. The model mesh contained 80x80 elements. Extracellular unipolar EGM were computed at each node of the isotropic grid as previously described using[1]:

$$u_{i,j}(t) = c \sum_{k,l=0}^{k,l=N} \frac{\vec{r}_{k,l} \nabla v_{k,l}(t)}{{r^3}_{k,l}}$$

where $u_{i,j}$ is the unipolar voltage at node $(i,j)$, $v_{k,l}$ the transmembrane voltage at node $(k,l)$ and $\vec{r}_{k,l}$ the distance between those nodes. A scaling coefficient is represented by $c$, which is assumed to equal 1 for simplicity.

Bipolar electrograms were computed as a difference of unipolar electrograms taken at a fixed spatial distance[2]:

$$w_{i,j}(t) = u_{i,j}(t) - u_{i+s,j}(t)$$

where $w_{i,j}$ is the bipolar voltage at node $(i, j)$ and $s$ represents the inter-electrode spacing (set to 3 in out study).

### S3 – Cleaning, Filtering, Phase Mapping and PS Detection

#### QRS Subtraction

A template subtraction method was used to remove far field ventricular depolarisation as described previously[3], following baseline correction of each epoch. Specifically, the template subtraction method identifies fiducial points for ventricular complexes using the QRS detection algorithm by Pan and Tompkins [4]. An average or 'median' ventricular complex was then constructed by aligning the detected ventricular complexes at their respective fiducial points, and performing a median operation of the matching points in all complexes [3]. Subtracting the median complex from each ventricular complex resulted in QRS subtracted electrograms.

#### Hilbert Transform Phase Reconstruction and PS Detection Methods

The instantaneous phase for each electrogram was reconstructed by applying the Hilbert transform on the cleaned and sinusoidally reconstructed signal [5, 6]. Mathematically, this can be given as follows:

$$\phi(t) = arctan\left(\frac{-u(t) - u*}{H(u)(t) - u*}\right)$$

where $u*$ sets the origin of the phase plane with respect to the phase that is computed [6].

#### PS Detection Methods

In this study, we use two approaches to detect PS: i) the classical line integral approach, and ii) the double ring approach. The classical PS detection approach is based on the line integral of the phase gradient [7]:

$$\oint_c \nabla\varphi d\vec{l}$$

where $\nabla$ is the spatial derivative, c is a closed loop surrounding a given point and ϕ is a phase map [7, 8]. Because the integral has to be discretised and approximated, c becomes the ring of electrode that encircles a point that is tested for the presence of a PS. Theoretically, the integral should result in ± 2 $\pi$ only at phase singularity points.

The double ring method is a modification of this, and consists of an inner 2 x 2 grid, enclosed by an outer 4 x 4 ring of electrodes. In a small 2 x 2 ring, PS detection translates to the presence of just a single phase difference greater than $\pi$, which is half of the full $2\pi$ cycle. As this may encompass more than one wave and cause artificial phase transitions, the second ring is added to ensure a PS is only identified if it is simultaneously detected by both the inner and outer ring [8]. This results in greater noise insensitivity.

#### S4 – Renewal Process Models

We model PS destruction and formation as renewal processes. For PS destruction, we measured the waiting times for an existing PS to be destroyed, and for PS formation, we studied the waiting times between the creation of new phase singularities. PS lifetime and inter-formation event times are random variables generated according to an exponential distribution, under the rate parameter λ. The probability density function for the all PS lifetimes or inter-formation times is thus given by:

$$f(t) = \begin{Bmatrix} 0 & t < 0 \\ \lambda e^{-\lambda t} & t \geq 0 \end{Bmatrix}$$

where $t$ is time, and $\lambda$ the PS destruction or formation rate.

#### Non-linear Least Squares Data Fitting

Parameter fitting to the exponential distribution was first completed using non-linear least squares on the probability distribution function (PDF) for each case. PDFs were generated by binning PS lifetime and inter-formation time data to form normalized histograms. Data fitting was performed in Matlab, and the fitting options selected include the Trust-Region Algorithm, with minimum difference change 1.0E-8, and maximum iterations set to 400. The r-squared ($R^2$) value was used to determine the adequacy of fit.

#### Maximum Likelihood Data Fitting

As a comparison to an alternative data fitting approach, PS data was also fitted using maximum likelihood. For continuous data, a histogram can throwaway information and fitting dependent on the choice of bin edges and bin widths. Maximum likelihood does not suffer from these problems, and as such was used as a comparative approach. Data fitting was performed in Matlab, using the probability density function (PDF) of each case. The chi-squared ($\chi 2$) good-ness of fit test was used to assess the adequacy of fit, which uses the null hypothesis that the data evaluated comes from an exponential distribution.

#### Data Fitting Results

To assess the adequacy of fit, we evaluated the $R^2$, sum of squares error (SSE), degrees of freedom error (DFE) and the standard error (RMSE) using non-linear least squares fitting. For the maximum likelihood approach, we determined the chi-squared ($\chi 2$) goodness-of-fit p-value, with $p < 0.05$ indicating that the null hypothesis is rejected and the data does not fit with the tested distribution

**Table 1- Destruction Process in Human Persistent AF**

| CASE | λ | 1/λ (μ) | R^2 | SSE | DFE | RMSE | $\chi^2$ p-valu |
|---|---|---|---|---|---|---|---|
| 1 | -0.0410 | -24.3780 | 0.9794 | 0.0009 | 155 | 0.0024 | 0.14 |
| 2 | -0.0358 | -27.9177 | 0.9023 | 0.0013 | 167 | 0.0028 | 0.92 |
| 3 | -0.0362 | -27.6473 | 0.9284 | 0.0005 | 216 | 0.0015 | 0.21 |
| 4 | -0.0448 | -22.3211 | 0.9100 | 0.0007 | 186 | 0.0019 | 0.32 |
| 5 | -0.0451 | -22.1816 | 0.9180 | 0.0006 | 169 | 0.0019 | 0.08 |
| 6 | -0.0511 | -19.5678 | 0.9554 | 0.0004 | 171 | 0.0016 | 0.09 |
| 7 | -0.0523 | -19.1335 | 0.9215 | 0.0007 | 175 | 0.0019 | 0.31 |
| 8 | -0.0369 | -27.0865 | 0.9484 | 0.0004 | 214 | 0.0013 | 0.45 |
| 9 | -0.0455 | -21.9864 | 0.9663 | 0.0003 | 219 | 0.0011 | 0.24 |
| 10 | -0.0753 | -13.2727 | 0.9639 | 0.0013 | 129 | 0.0031 | 0.92 |
| 11 | -0.0522 | -19.1610 | 0.9878 | 0.0010 | 126 | 0.0028 | 0.27 |
| 12 | -0.0681 | -14.6820 | 0.9579 | 0.0014 | 142 | 0.0032 | 0.77 |
| 13 | -0.0506 | -19.7622 | 0.9438 | 0.0019 | 95 | 0.0045 | 0.19 |
| 14 | -0.0388 | -25.7533 | 0.9444 | 0.0017 | 132 | 0.0036 | 0.29 |
| 15 | -0.0423 | -23.6537 | 0.9689 | 0.0002 | 225 | 0.0011 | 0.09 |
| 16 | -0.0370 | -27.0051 | 0.9091 | 0.0006 | 186 | 0.0019 | 0.58 |

| | | | | | | | |
|---|---|---|---|---|---|---|---|
| **17** | -0.0311 | -32.1554 | 0.9977 | 0.0015 | 108 | 0.0037 | 0.68 |
| **18** | -0.0322 | -31.0795 | 0.9364 | 0.0011 | 110 | 0.0032 | 0.55 |
| **19** | -0.0507 | -19.7213 | 0.9373 | 0.0006 | 187 | 0.0017 | 0.43 |
| **20** | -0.0381 | -26.2146 | 0.9083 | 0.0006 | 177 | 0.0019 | 0.64 |
| **21** | -0.0431 | -23.2038 | 0.9900 | 0.0007 | 159 | 0.0022 | 0.65 |
| **22** | -0.0595 | -16.8024 | 0.9852 | 0.0015 | 121 | 0.0035 | 0.68 |
| **23** | -0.0381 | -26.2131 | 0.9822 | 0.0001 | 280 | 0.0007 | 0.64 |
| **24** | -0.0390 | -25.6334 | 0.9165 | 0.0006 | 181 | 0.0018 | 0.95 |
| **25** | -0.0435 | -23.0031 | 0.9196 | 0.0006 | 167 | 0.0019 | 0.21 |
| **26** | -0.0316 | -31.6781 | 0.9152 | 0.0005 | 225 | 0.0016 | 0.71 |
| **27** | -0.0339 | -29.4629 | 0.9839 | 0.0009 | 164 | 0.0024 | 0.24 |
| **28** | -0.0436 | -22.9554 | 0.8927 | 0.0012 | 112 | 0.0033 | 0.12 |
| **29** | -0.0504 | -19.8610 | 0.9335 | 0.0006 | 167 | 0.0019 | 0.61 |
| **30** | -0.0556 | -17.9914 | 0.9698 | 0.0012 | 119 | 0.0031 | 0.45 |
| **31** | -0.0637 | -15.7036 | 0.9166 | 0.0009 | 129 | 0.0026 | 0.46 |
| **32** | -0.0497 | -20.1090 | 0.9147 | 0.0008 | 160 | 0.0022 | 0.66 |
| **33** | -0.0552 | -18.1207 | 0.9096 | 0.0008 | 146 | 0.0024 | 0.77 |
| **34** | -0.0429 | -23.3163 | 0.9876 | 0.0008 | 159 | 0.0023 | 0.35 |
| **35** | -0.0530 | -18.8568 | 0.9190 | 0.0010 | 153 | 0.0025 | 0.66 |
| **36** | -0.0362 | -27.5905 | 0.9900 | 0.0006 | 179 | 0.0019 | 0.42 |

| | | | | | | | |
|---|---|---|---|---|---|---|---|
| **37** | -0.0455 | -21.9873 | 0.9170 | 0.0007 | 187 | 0.0019 | 0.84 |
| **38** | -0.0452 | -22.1229 | 0.9060 | 0.0007 | 178 | 0.0020 | 0.83 |
| **39** | -0.0435 | -22.9993 | 0.9729 | 0.0010 | 172 | 0.0024 | 0.26 |
| **40** | -0.0433 | -23.1048 | 0.9154 | 0.0007 | 160 | 0.0020 | 0.61 |
| **41** | -0.0489 | -20.4399 | 0.9141 | 0.0007 | 191 | 0.0020 | 0.58 |
| **42** | -0.0465 | -21.5199 | 0.9090 | 0.0008 | 165 | 0.0022 | 0.54 |
| **43** | -0.0620 | -16.1247 | 0.9873 | 0.0011 | 149 | 0.0027 | 0.87 |

**Table 2- Formation Process in Human Persistent AF**

| **CASE** | **λ** | **1/λ (μ)** | **R^2** | **SSE** | **DFE** | **RMSE** | **$\chi^2$ p-value** |
|---|---|---|---|---|---|---|---|
| **1** | -0.0424 | -23.601 | 0.9798 | 0.0009 | 260 | 0.0019 | 0.26 |
| **2** | -0.0540 | -18.526 | 0.9291 | 0.0011 | 187 | 0.0025 | 0.32 |
| **3** | -0.0377 | -26.529 | 0.9545 | 0.0007 | 343 | 0.0014 | 0.12 |
| **4** | -0.0496 | -20.145 | 0.9001 | 0.0006 | 226 | 0.0016 | 0.94 |
| **5** | -0.0523 | -19.116 | 0.9476 | 0.0010 | 273 | 0.0019 | 0.65 |
| **6** | -0.0442 | -22.615 | 0.9041 | 0.0005 | 378 | 0.0011 | 0.48 |
| **7** | -0.0516 | -19.381 | 0.9599 | 0.0009 | 227 | 0.0020 | 0.64 |
| **8** | -0.0537 | -18.623 | 0.9381 | 0.0005 | 260 | 0.0013 | 0.54 |
| **9** | -0.0525 | -19.040 | 0.9411 | 0.0004 | 307 | 0.0012 | 0.65 |
| **10** | -0.0575 | -17.405 | 0.9168 | 0.0009 | 374 | 0.0016 | 0.54 |

| | | | | | | | |
|---|---|---|---|---|---|---|---|
| **11** | -0.0401 | -24.912 | 0.9327 | 0.0011 | 292 | 0.0020 | 0.72 |
| **12** | -0.0412 | -24.285 | 0.9566 | 0.0010 | 310 | 0.0018 | 0.52 |
| **13** | -0.0596 | -16.784 | 0.9302 | 0.0017 | 203 | 0.0029 | 0.99 |
| **14** | -0.0379 | -26.377 | 0.9576 | 0.0010 | 195 | 0.0022 | 0.22 |
| **15** | -0.0387 | -25.856 | 0.9296 | 0.0003 | 361 | 0.0010 | 0.11 |
| **16** | -0.0385 | -26.004 | 0.9425 | 0.0005 | 274 | 0.0014 | 0.11 |
| **17** | -0.0636 | -15.735 | 0.6165 | 0.0017 | 338 | 0.0023 | 0.06 |
| **18** | -0.0753 | -13.279 | 0.9074 | 0.0011 | 367 | 0.0017 | 0.40 |
| **19** | -0.0207 | -48.242 | 0.9071 | 0.0039 | 20 | 0.0139 | 0.45 |
| **20** | -0.0483 | -20.690 | 0.9793 | 0.0007 | 246 | 0.0017 | 0.37 |
| **21** | -0.0515 | -19.431 | 0.9661 | 0.0008 | 266 | 0.0017 | 0.76 |
| **22** | -0.0397 | -25.165 | 0.9647 | 0.0012 | 333 | 0.0019 | 0.63 |
| **23** | -0.0207 | -48.265 | 0.9501 | 0.0057 | 50 | 0.0107 | 0.77 |
| **24** | -0.0428 | -23.362 | 0.9329 | 0.0006 | 269 | 0.0015 | 0.93 |
| **25** | -0.0445 | -22.487 | 0.9374 | 0.0006 | 267 | 0.0015 | 0.97 |
| **26** | -0.0458 | -21.812 | 0.9415 | 0.0006 | 223 | 0.0017 | 0.19 |
| **27** | -0.0479 | -20.872 | 0.9184 | 0.0011 | 205 | 0.0023 | 0.14 |
| **28** | -0.0331 | -30.232 | 0.9157 | 0.0019 | 216 | 0.0030 | 0.70 |
| **29** | -0.0566 | -17.676 | 0.9889 | 0.0007 | 306 | 0.0015 | 0.09 |
| **30** | -0.0589 | -16.974 | 0.9689 | 0.0019 | 196 | 0.0031 | 0.53 |

| | | | | | | | |
|---|---|---|---|---|---|---|---|
| **31** | -0.0566 | -17.675 | 0.9468 | 0.0009 | 281 | 0.0018 | 0.53 |
| **32** | -0.0551 | -18.140 | 0.9899 | 0.0007 | 284 | 0.0015 | 0.86 |
| **33** | -0.0476 | -21.004 | 0.9074 | 0.0010 | 323 | 0.0017 | 0.48 |
| **34** | -0.0226 | -44.284 | 0.9577 | 0.0008 | 310 | 0.0016 | 0.39 |
| **35** | -0.0423 | -23.613 | 0.9782 | 0.0009 | 319 | 0.0017 | 0.67 |
| **36** | -0.0384 | -26.026 | 0.9870 | 0.0005 | 278 | 0.0014 | 0.74 |
| **37** | -0.0573 | -17.463 | 0.9883 | 0.0008 | 287 | 0.0017 | 0.52 |
| **38** | -0.0545 | -18.355 | 0.9620 | 0.0009 | 301 | 0.0017 | 0.35 |
| **39** | -0.0339 | -29.468 | 0.9681 | 0.0009 | 292 | 0.0017 | 0.15 |
| **40** | -0.0489 | -20.449 | 0.9605 | 0.0008 | 293 | 0.0016 | 0.59 |
| **41** | -0.0457 | -21.871 | 0.9847 | 0.0005 | 267 | 0.0014 | 0.26 |
| **42** | -0.0436 | -22.923 | 0.9736 | 0.0006 | 291 | 0.0015 | 0.08 |
| **43** | -0.0392 | -25.510 | 0.9227 | 0.0007 | 313 | 0.0015 | 0.75 |

**Table 3- Destruction Process in Rat AF**

| CASE | Λ | μ | R^2 | SSE | DFE | RSME |
|---|---|---|---|---|---|---|
| **1** | -0.39464 | -2.53397 | 0.920077 | 0.001655 | 19 | 0.015156 |
| **2** | -0.32009 | -3.12408 | 0.906601 | 0.001161 | 24 | 0.014547 |
| **3** | -0.45308 | -2.2071 | 0.910141 | 0.001666 | 25 | 0.014645 |
| **4** | | | NO PS | | | |

**Table 4- Destruction Process in Rat VF**

| CASE | Λ | μ | R^2 | SSE | DFE | RSME |
|---|---|---|---|---|---|---|
| **1** | -0.616976139 | -1.62081 | 0.925768 | 0.005767 | 16 | 0.018985 |
| **2** | -0.762581075 | -1.31134 | 0.965581 | 0.003513 | 23 | 0.012359 |
| **3** | -0.499943945 | -2.00022 | 0.935486 | 0.004449 | 27 | 0.012837 |
| **4** | -0.392332499 | -2.54886 | 0.952071 | 0.025498 | 18 | 0.037637 |
| **5** | -0.386703801 | -2.58596 | 0.915747 | 0.008958 | 17 | 0.022955 |
| **6** | -0.393236373 | -2.543 | 0.949446 | 0.007835 | 18 | 0.020863 |
| **7** | -0.571397065 | -1.7501 | 0.92278 | 0.020703 | 11 | 0.043383 |
| **8** | -0.791982816 | -1.26265 | 0.938596 | 0.011686 | 12 | 0.031207 |
| **9** | -0.099075603 | -10.0933 | 0.921393 | 0.021087 | 13 | 0.040275 |
| **10** | -0.263687202 | -3.79237 | 0.965384 | 0.01155 | 18 | 0.025331 |
| **11** | -0.251567598 | -3.97507 | 0.96888 | 0.009258 | 19 | 0.022075 |

**Table 5- Destruction Process in Tachypaced Sheep AF**

| CASE | λ | 1/λ () | R^2 | SSE | DFE | RMSE | $\chi^2$ p-value |
|---|---|---|---|---|---|---|---|
| **1** | -0.0579 | -17.2641 | 0.9768 | 0.0013 | 152 | 0.00288 | 0.77 |
| **2** | -0.0432 | -23.1674 | 0.9256 | 0.0006 | 173 | 0.00182 | 0.40 |
| **3** | -0.0409 | -24.4553 | 0.9899 | 0.0007 | 176 | 0.00204 | 0.81 |
| **4** | -0.0438 | -22.8181 | 0.9373 | 0.0005 | 184 | 0.00166 | 0.76 |
| **5** | -0.0461 | -21.6732 | 0.9067 | 0.0008 | 185 | 0.00205 | 0.38 |
| **6** | -0.0448 | -22.3027 | 0.9389 | 0.0005 | 183 | 0.00164 | 0.22 |
| **7** | -0.0503 | -19.8749 | 0.9169 | 0.0015 | 122 | 0.00346 | 0.79 |

| | | | | | | | |
|---|---|---|---|---|---|---|---|
| **8** | -0.0515 | -19.4243 | 0.9267 | 0.0016 | 137 | 0.00339 | 0.95 |
| **9** | -0.0459 | -21.7629 | 0.9053 | 0.0015 | 144 | 0.00324 | 0.33 |
| **10** | -0.0497 | -20.1324 | 0.9405 | 0.0013 | 164 | 0.00285 | 0.67 |
| **11** | -0.0425 | -23.5366 | 0.9556 | 0.0012 | 157 | 0.00271 | 0.44 |
| **12** | -0.0454 | -22.041 | 0.9375 | 0.0012 | 165 | 0.00271 | 0.83 |
| **13** | -0.0419 | -23.8595 | 0.9546 | 0.0011 | 183 | 0.00242 | 0.77 |
| **14** | -0.0360 | -27.8098 | 0.9551 | 0.0009 | 192 | 0.00218 | 0.17 |
| **15** | -0.0411 | -24.3285 | 0.9793 | 0.0014 | 200 | 0.00265 | 0.86 |
| **16** | -0.0368 | -27.1747 | 0.9763 | 0.0008 | 159 | 0.00227 | 0.99 |
| **17** | -0.0368 | -27.1681 | 0.9402 | 0.0011 | 156 | 0.00263 | 0.51 |
| **18** | -0.0342 | -29.229 | 0.9550 | 0.0009 | 165 | 0.00233 | 0.88 |

**Table 6- Formation Process in Tachypaced Sheep AF**

| CASE | λ | 1/λ () | R^2 | SSE | DFE | RMSE | $\chi^2$ p-value |
|---|---|---|---|---|---|---|---|
| **1** | -0.0473 | -21.1452 | 0.9333 | 0.0010 | 233 | 0.0020 | 0.24 |
| **2** | -0.0487 | -20.5526 | 0.9851 | 0.0007 | 197 | 0.0019 | 0.44 |
| **3** | -0.0459 | -21.7789 | 0.9662 | 0.0007 | 200 | 0.0019 | 0.69 |
| **4** | -0.0552 | -18.1025 | 0.9297 | 0.0005 | 176 | 0.0017 | 0.36 |
| **5** | -0.0552 | -18.1040 | 0.9135 | 0.0006 | 179 | 0.0019 | 0.74 |
| **6** | -0.0550 | -18.1693 | 0.9889 | 0.0008 | 167 | 0.0022 | 0.39 |

| | | | | | | | |
|---|---|---|---|---|---|---|---|
| **7** | -0.0367 | -27.2372 | 0.9264 | 0.0014 | 250 | 0.0024 | 0.68 |
| **8** | -0.0491 | -20.3860 | 0.9733 | 0.0012 | 257 | 0.0022 | 0.70 |
| **9** | -0.0442 | -22.6277 | 0.9121 | 0.0014 | 251 | 0.0023 | 0.44 |
| **10** | -0.0456 | -21.9182 | 0.9106 | 0.0010 | 237 | 0.0021 | 0.09 |
| **11** | -0.0467 | -21.4319 | 0.9203 | 0.0010 | 249 | 0.0020 | 0.33 |
| **12** | -0.0407 | -24.5926 | 0.9740 | 0.0010 | 250 | 0.0020 | 0.42 |
| **13** | -0.0434 | -23.0544 | 0.9788 | 0.0011 | 235 | 0.0022 | 0.27 |
| **14** | -0.0458 | -21.8345 | 0.9385 | 0.0008 | 247 | 0.0018 | 0.20 |
| **15** | -0.0456 | -21.9338 | 0.9455 | 0.0014 | 238 | 0.0024 | 0.82 |
| **16** | -0.0400 | -24.9770 | 0.9153 | 0.0009 | 228 | 0.0020 | 0.43 |
| **17** | -0.0372 | -26.8711 | 0.9783 | 0.0010 | 231 | 0.0020 | 0.89 |
| **18** | -0.0381 | -26.2718 | 0.9788 | 0.0010 | 233 | 0.0020 | 0.39 |

**Table 7- Formation Process in Rat AF**

| CASE | Λ | M | R^2 | SSE | DFE | RSME |
|---|---|---|---|---|---|---|
| **1** | 0.419942 | 2.381283 | 0.942991 | 0.001655 | 17 | 0.005932 |
| **2** | 0.243807 | 4.101607 | 0.931647 | 0.001161 | 19 | 0.00598 |
| **3** | 0.315868 | 3.165879 | 0.94344 | 0.001666 | 8 | 0.00343 |
| **4** | | | NO PS | | | |

**Table 8- Formation Process in Rat VF**

| CASE | Λ | M | R^2 | SSE | DFE | RSME |
|---|---|---|---|---|---|---|
| **1** | 0.654083648 | 1.528856 | 0.911355 | 0.001497 | 30 | 0.007063 |
| **2** | 0.630159282 | 1.5869 | 0.970007 | 0.002665 | 26 | 0.010125 |
| **3** | 0.130750854 | 7.648134 | 0.939085 | 0.001106 | 20 | 0.007438 |
| **4** | 0.465554173 | 2.147978 | 0.960541 | 0.001834 | 39 | 0.006858 |
| **5** | 0.359369029 | 2.782655 | 0.904757 | 0.003788 | 30 | 0.011236 |
| **6** | 0.195596497 | 5.112566 | 0.95903 | 0.008241 | 33 | 0.015803 |
| **7** | 0.219044658 | 4.565279 | 0.986648 | 0.004465 | 29 | 0.012409 |
| **8** | 0.113454978 | 8.814069 | 0.995203 | 0.014203 | 22 | 0.025408 |
| **9** | 0.863465002 | 1.158125 | 0.917717 | 0.018902 | 20 | 0.030743 |
| **10** | 0.280973897 | 3.559049 | 0.987404 | 0.004937 | 33 | 0.012231 |
| **11** | 0.310418765 | 3.221455 | 0.911724 | 0.002504 | 37 | 0.008226 |

#### S5 – Effect of Electrode Permutation

Electrode permutation resulted in an increased number of PS detections using the line integral approach (un-permuted count: 330 ± 6.4 PS; permuted count: 415 ± 4.7 PS). Contrastingly, the double ring approach decreased the number of PS detections (un-permuted count: 261 ± 1.9 PS; permuted count: 257 ± 2.0 PS), confirming its noise insensitivity.

Permuted data consistently yielded lower observed and MaxEnt $\lambda$ for both PS detection methods, with mean $\lambda$ equalling 2.8 %/ms for unshuffled data, and dropping to 1.2 %s PS/s for shuffled data using the line integral approach. Results were similar for the double ring approach (un-permuted- $\lambda$: 4.4 %m/s; permuted $\lambda$: 2.3 %/ms). A one sample t-test verified consistent statistical significance between permuted and un-permuted $\lambda$ ($p < 0.001$ for all cases).

#### S6 – Comparison between Double Ring and Line Integral PS Detection Methods

In this study, we used the two PS detection methods to construct a PS look-up table indexing the time of onset and location for each new PS (as previously described in S2). This allowed us to determine PS lifetime data and PS inter-formation event times, and therefore PS destruction and creation rates. Using both PS detection methods, the predicted and observed distributions of PS lifetimes and inter-formation event times were consistently exponential as shown in Figure 1. This suggests that the underlying PS destruction process is truly exponential irrespective of the PS detection method used.

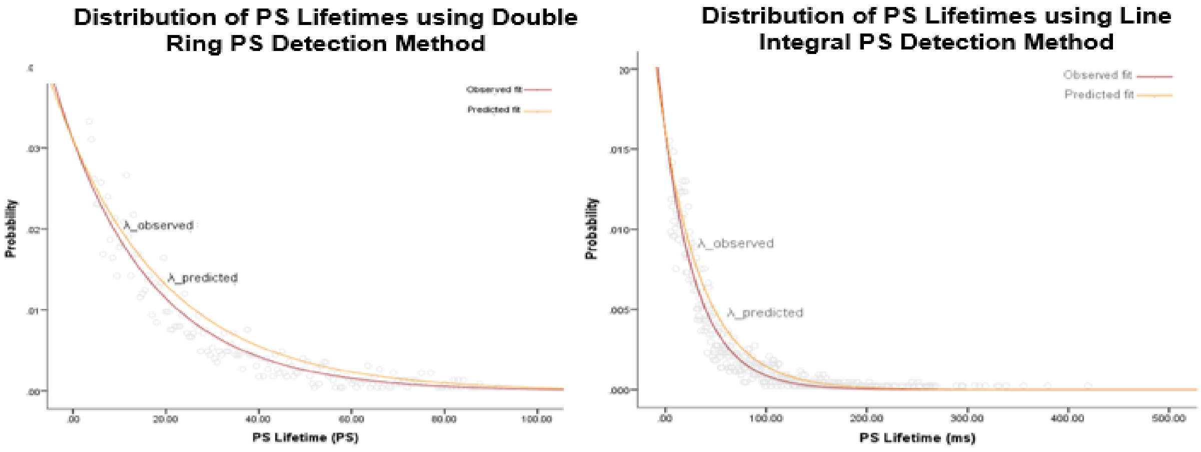


*Figure 1: Comparison of PS Lifetime Curves using Different PS Detection Methods*

The left and right hand figures show the PS lifetime curve (with $\lambda_{obs}$ and $\lambda_{d\text{-}pred}$, which was computed using principles of MaxEnt) using two different PS detection methods: i) the double ring approach (left) and ii) classical line integral approach (right). Both methods yield an exponential distribution of both the observed and predicted curves, suggesting that the underlying PS destruction process is truly exponential irrespective of the PS detection method used.

#### S7 – A Simple Step by Step Guide for λ Estimation Using MaxEnt Principles and Lagrange Multipliers

The general principle behind MaxEnt is that the optimal choice for the predicted probability distribution of a given data set is one with the largest amount of uncertainty or 'entropy', e.g. the choice that is the most purely random containing the least number of assumptions, while still meeting the known constraints. To achieve this, we can maximize the entropy function subject to our constraints using the Lagrange multiplier method:

$$\frac{\partial f}{\partial x} = \sum_{i=1}^{N} \lambda_i \frac{\partial g_x}{\partial x}$$

where $N$ is the number of constraints, $g_x(x)$ are the constraints, $\lambda_i$ the Lagrange multiplier, and $f(x)$ is the entropy function:

$$f(x) = -\sum_{x=0}^{\infty} (p_x)\log(p_x)$$

In general terms, the Lagrange multiplier itself is a variable that allows the directions and gradients of $f(x)$ and $g(x)$ to be compared, despite their respective magnitudes not necessarily being equal. This allows the Lagrange multiplier method to find the minima and maxima of a function subject to constraints without having to explicitly solve the conditions.

For a waiting-time process, the lowest moment of the sample distribution, the sample mean, is applied as the constraint to achieve the most unbiased probability distribution ($P_{MaxEnt}$). In this study specifically, the sample mean is equal to the mean of PS lifetimes or PS inter-formation event times for each AF epoch. As probabilities must always sum to 1, a normalization constraint must also be applied. Therefore, we must find $P_{MaxEnt}$ such that:

1. $P_{MaxEnt}$ satisfies the constraint on the average sample mean
2. $P_{MaxEnt}$ satisfies the normalization constraint
3. $P_{MaxEnt}$ has the maximum entropy

Mathematically, such a distribution can be found by setting the gradient of the function to equal a linear combination of the gradients of the constraints:

$$\frac{\partial f}{\partial p_x} = \lambda_1 \frac{\partial g_1}{\partial p_x} + \lambda_2 \frac{\partial g_2}{\partial p_x} \quad \textbf{(1)}$$

where $f(x)$ is the entropy function and $g_1(x)$ is the expectation of $X$ (sample mean)

$$\begin{aligned} g_1(x) &= E[X] \\ &= \sum x.p_x \\ &= \langle \mathrm{w} \rangle \end{aligned} \quad \textbf{(2)}$$

Where <w> can be estimated by the sample mean.

Further:

$$\therefore \frac{\partial g_1}{\partial p_x} = x$$

and $g_2(x)$ the normalization constraint:

$$g_2(x) = \sum p_x \qquad (3)$$

$$= 1$$

$$\therefore \frac{\partial g_2}{\partial p_x} = 1$$

Therefore, solving equation (1) gives the $P_{MaxEnt}$ of PS lifetimes and PS inter-formation event times. This can be achieved by following the steps as outlined below:

To find ∂f/∂px, we use the chain rule:

$$f(x) = -\sum_{x=0}^{\infty}(p_x)\log(p_x)$$

$$\frac{\partial f}{\partial px} = -((\log(p_x) + (p_x).\frac{1}{p_x})$$

$$= -(logp_x + 1)$$

Substituting into equation 1

$$-(log(p_x) + 1) = \lambda_1(x) + \lambda_2(1) \qquad (4)$$

$$\therefore -(\log(p_x) + 1) = \lambda_1(x) + \lambda_2 \qquad (5)$$

1. Solving for $p_x$:

$$p_x = e^{-1-\lambda_1 x-\lambda_2} \qquad (6)$$

$$= e^{-\lambda_1 x}.e^{-(\lambda_2+1)}$$

$$p_x = \frac{e^{-\lambda_1 x}}{Z} \qquad (7)$$

$$\text{where Z} = e^{1+\lambda_2} \qquad (8)$$

As we can see, the functional form of $P_{MaxEnt}$ gives an exponential distribution. As a result, solving for $\lambda_1$will give the predicted exponential decay constant of the distribution, and in turn the predicted PS destruction and formation rate. This was used to compare with the observed $\lambda$ from the experimental data. To solve for $\lambda_1$, we must solve for Z:

2. Plugging in constraint $g_2$ (normalization constraint) into equation 7 gives:

$$\frac{1}{Z}\sum_{0}^{\infty} e^{-\lambda_1 x} = 1 \quad \textbf{(9)}$$

$$\therefore Z = \sum_{0}^{\infty} e^{-\lambda_1 x} \quad \textbf{(10)}$$

Using geometric series expansion

**(11)**

$$= 1 + e^{-\lambda_1 +} e^{-2\lambda_1 + \cdots}$$

$$Z = \frac{1}{1 - e^{-\lambda_1}}$$

$$\therefore \frac{\partial Z}{\partial \lambda_1} = \frac{e^{-\lambda_1}}{(1 - e^{-\lambda_1})^2}$$

3. Plugging in constraint and $g_1$ (sample mean constraint) into equation 7 gives:

$$g_1(x) \; = \sum x.p_x \quad \textbf{(12)}$$

$$= \sum x \frac{e^{-\lambda_1 x}}{Z}$$

$$= \frac{1}{Z}\sum_{0}^{\infty} x e^{-\lambda_1 x}$$

$$= < w >$$

4. To help solve for equation (12), we take the derivative of Z (equation 10):

$$\frac{\partial Z}{\partial \lambda_1} = -\sum_{0}^{\infty} x e^{-\lambda_1 x} \quad \textbf{(13)}$$

5. Therefore we can re-write the first constraint equation (12) as:

$$g_1(x) = \frac{1}{Z}\sum_{0}^{\infty} x e^{-\lambda_1 x} \quad \textbf{(14)}$$

$$= \frac{1}{Z} * \frac{-\partial Z}{\partial \lambda_1} \quad \textbf{(15)}$$

$$= < w >$$

$$\therefore \frac{1}{Z}\frac{\partial Z}{\partial \lambda_1} = \frac{\left(1 - e^{-\lambda_1}\right)}{1} * \frac{e^{-\lambda_1}}{(1 - e^{-\lambda_1})^2}$$

$$= \frac{e^{-\lambda_1}}{1 - e^{-\lambda_1}}$$

$$= < w >$$

Solving for equation (15) will therefore give $\lambda_1$ and in turn the PS destruction or formation rate.

#### S8- Systematic Review of Literature for PS lifetime data

The search criteria was: ‘atrial fibrillation OR ventricular fibrillation AND rotor* OR phase singularit* OR spiral wave*.’ Ventricular fibrillation was also included as it was reasoned that a similar stochastic process could apply in VF, and a subgroup analysis was performed for these data. The criteria for inclusion were studies in humans or experimental model systems reporting histogram or probability distribution data for phase singularity lifetimes. Studies were excluded if they were reviews, editorials, or commentary. Candidate full-text studies were retrieved for secondary review, with final inclusion determined by consensus.

For each study, histograms of PS lifetime were digitized, with probabilities measured with digital calipers (DigitizeIt, Koln, Germany). λ observed was computed from the exponential decay constant of the digitized PDF. To cross validate, the λ MaxEnt was computed using the principle of Maximum entropy and Langrangian multipliers.

The results are shown in Table 1, with relevant panels shown in Figure 6 showing the distributional shape of PS lifetimes. The MaxEnt predicted λ and observed data λ showed a consistent association (Table 1) with parameters similar to found in our data

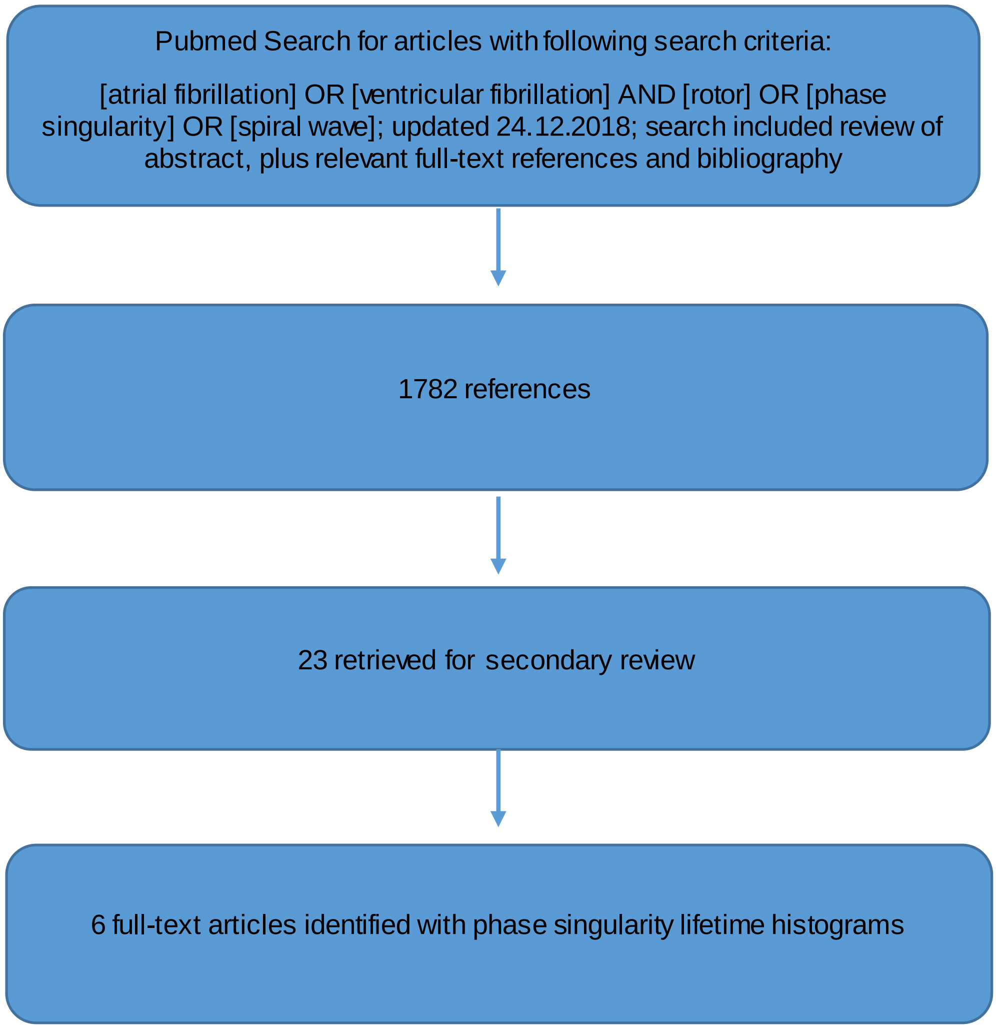

**TABLE 1– Systematic Review Results – Studies showing histogram or probability distribution for phase singularity lifetime in AF and VF 2000-2018.**

| YEAR | FIRST AUTHOR | PRINCIPAL INVESTIGATOR | JOURNAL | ARRHYTHMIA | MODEL SYSTEM | λ MaxEnt (%/ms)+ | λ observed (%/ms)++ |
|---|---|---|---|---|---|---|---|
| **2000** | Chen[24] | J.Jalifé, SUNY | Circ Res | VF | Sheep, optical mapping | 6.8%/ms | 6.1%/ms |
| **2000** | Chen[25] | Jalifé,SUNY | Cardiovasc Res | AF | Sheep, Optical mapping | 5.1%/ms | 5.2%/ms |
| **2006** | Kay[26] | J.Rogers, University of Alabama | Am J Phys Heart Circ Phys | VF | Porcine optical mapping | * | 2.7 %/ms |
| **2017** | Kuklik[22] | Schötten | IEEE | AF | Contact plaque unipolar electrograms from cardiac surgery | ** | 1.6%/ms |
| **2018** | Child[17] | J.Gill, | Circ AE | AF | Human,basket catheter during AF ablation | 2.7%/ms | 3.4%/ms |
| **2018** | Christoph[27] | S.Luther | Nature | VF | Optical mapping, electromechanical mapping | 2.1%/ms | 2.9%/ms |

+ Predicted λ is derived from expectation of PS lifetime using Lagrangian multipliers as shown in Methods

++ Observed λ is determined from nonlinear least squares fitted coefficient for the probability distribution for PS lifetime

*Estimated based on text that 0.996 of PS were <200ms, MaxEnt prediction unable be calculated as sample mean for PS lifetime because binning was too wide (200ms). PS –observed based on curve fit.

** MaxEnt prediction unable to calculate sample mean for PS lifetime as binning too wide (by rotation number) to allow estimation of sample mean lifetime; λ -observed based on fitting curve to histogram with estimated PS lifetime for 1 rotation of 160ms.

#### S9 – λ By Anatomical Chamber

For PS destruction, there were no statistically significant differences (P = 0.18) in PS destruction λ between the LA (4.5 %/ms (95%CI, 4.1, 4.8)) and RA (4.9 %/ms (95%CI, 4.3, 5.4), S3) in humans. Mean λ in sheep between LA (4.5 %/ms (95%CI, 4.02, 5.1)) and RA (4.22 %/ms (95%CI, 3.8, 4.6)) was also not statistically different between chambers (P=0.20) (S3).

For PS Formation, human λ for PS formation in the LA (4.9 %/ms (95%CI, 4.5, 5.4)) (4.1 %/ms (95%CI, 3.6, 4.6) (P=0.28)), which was not statistically significant. PS formation in the LA (LA: 4.5 %ms/s (95%CI, 5.1, 6.7) was also not different to the RA: 4.6 %ms/s (95%CI, 5.3, 6.9); P = 0.79) for sheep.

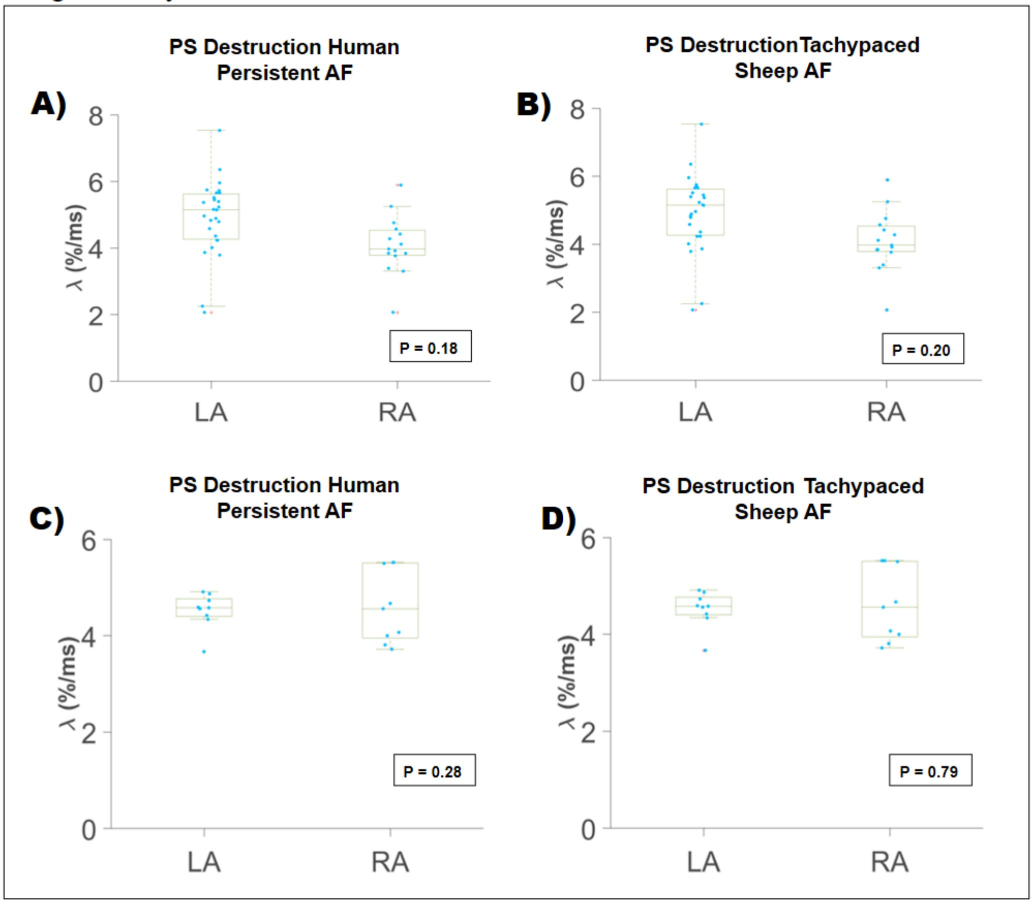


***Figure 2: λ by Anatomical Chamber***
**PS Destruction (2A-B):** For human persistent AF, there were no statistically significant differences (P = 0.18) in PS destruction λ between the LA and RA (**5A**). Tachypaced sheep AF also showed no differences between the LA and RA (P = 0.20) (**5B**). **PS Creation (5C-D):** No statistically significant inter-chamber differences were seen between λ in the LA and RA in persistent human AF (**5C**) (P = 0.28), or tachypaced sheep AF (0.79) (**5D**).

### S10 – Correlation between Experimental λ and MaxEnt Predicted λ

To confirm that the underlying data generating process responsible for PS formation and destruction are purely stochastic, we compared distributions and λ from the experimental data to those predicted by Maximum entropy (MaxEnt).

The MaxEnt predicted λ consistently highly correlated with λ from the experimental data.

Fig 7. MaxEnt Predicted and Fitted λ are Highly Correlated

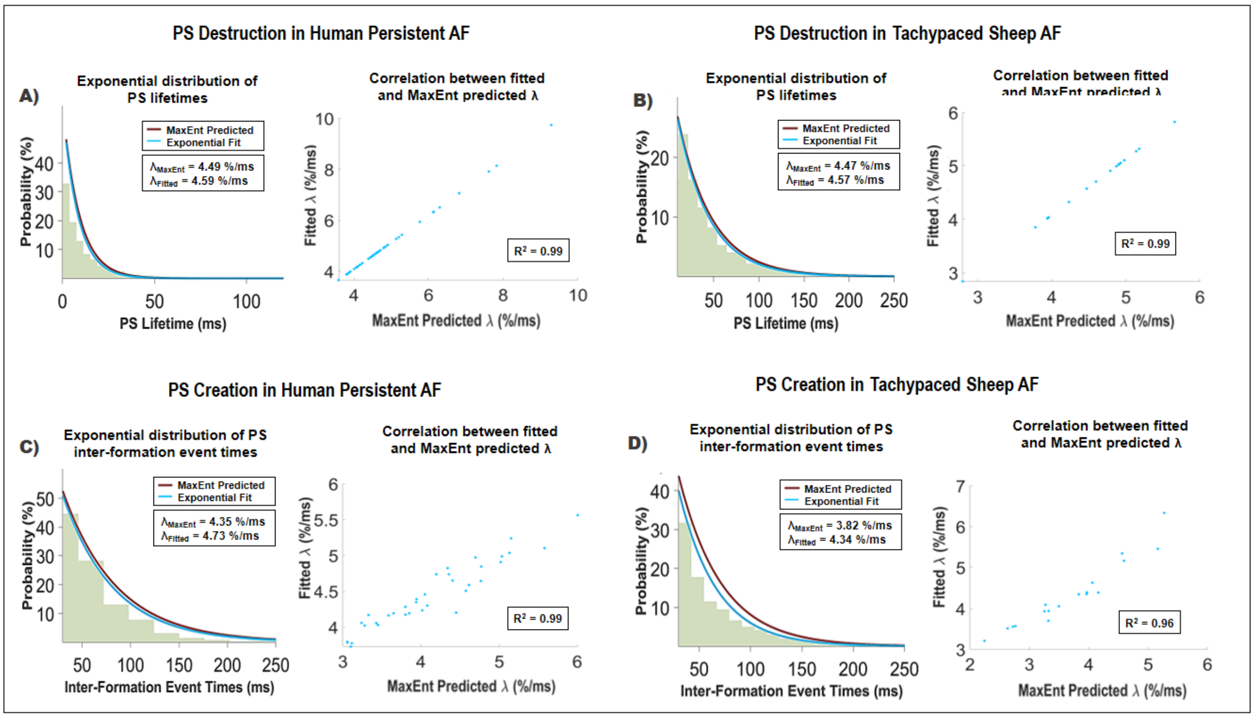


***Figure 3: MaxEnt Predicted and Fitted λ are Highly Correlated***

***PS Destruction in Human Persistent AF and Tachypaced Sheep AF (3A-B):*** PS lifetime distributions were exponential for both MaxEnt predicted and maximum likelihood fitted distributions (left). MaxEnt predicted and fitted were highly correlated (right), suggesting underlying data generating process is maximally random and stochastic (e.g. a Poisson renewal process).

***PS Creation in Human Persistent AF and Tachypaced Sheep AF (3C-D):*** PS inter-formation event time distributions were exponential for both MaxEnt predicted and maximum likelihood fitted distributions (left). MaxEnt predicted and fitted were highly correlated (right).

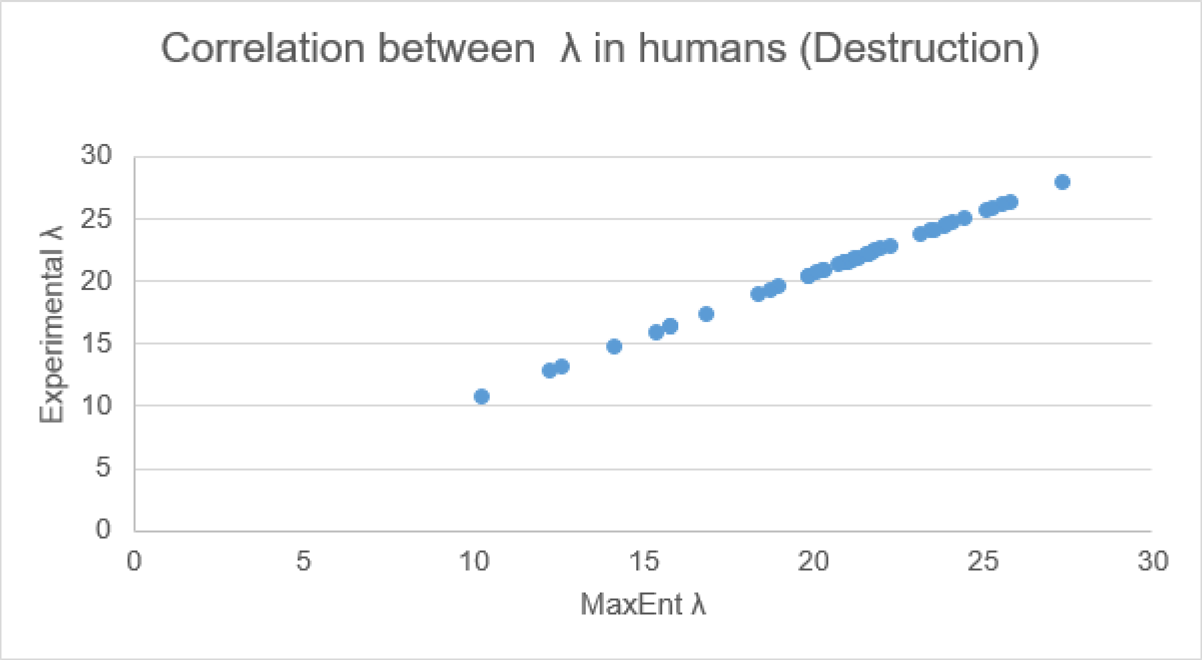
Correlation between λ in humans (Destruction)
Experimental λ
30
25
20
15
10
5
0
0
5
10
15
20
25
30
MaxEnt λ

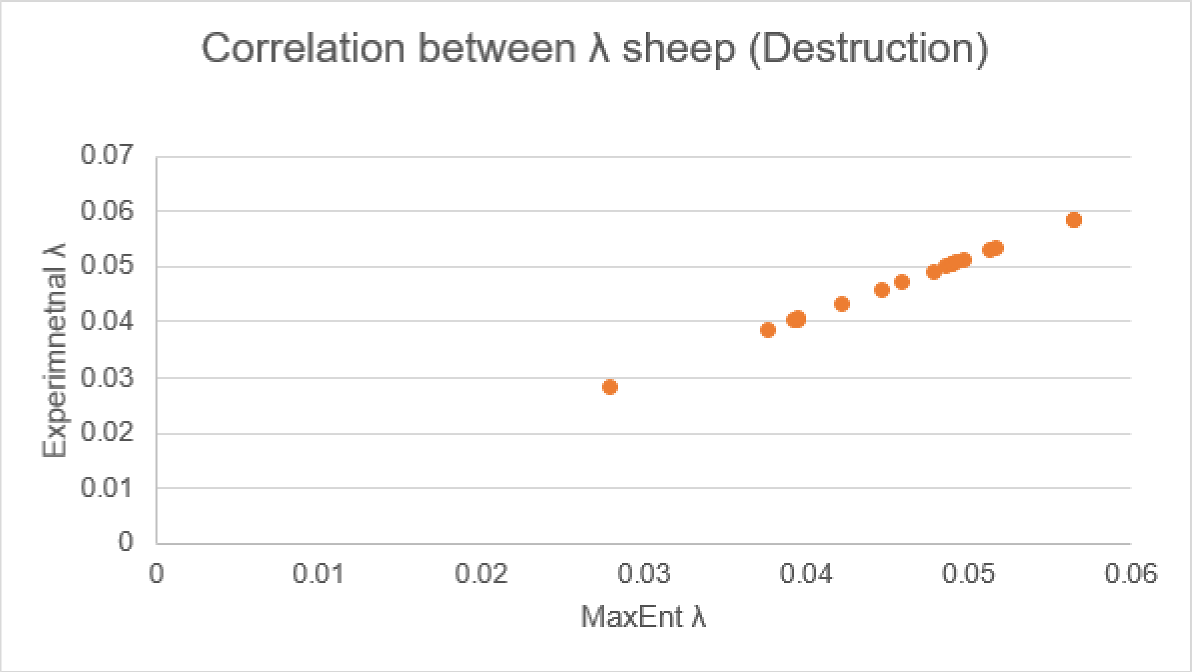
Correlation between λ sheep (Destruction)
Experimnetnal λ
0.07
0.06
0.05
0.04
0.03
0.02
0.01
0
0
0.01
0.02
0.03
0.04
0.05
0.06
MaxEnt λ

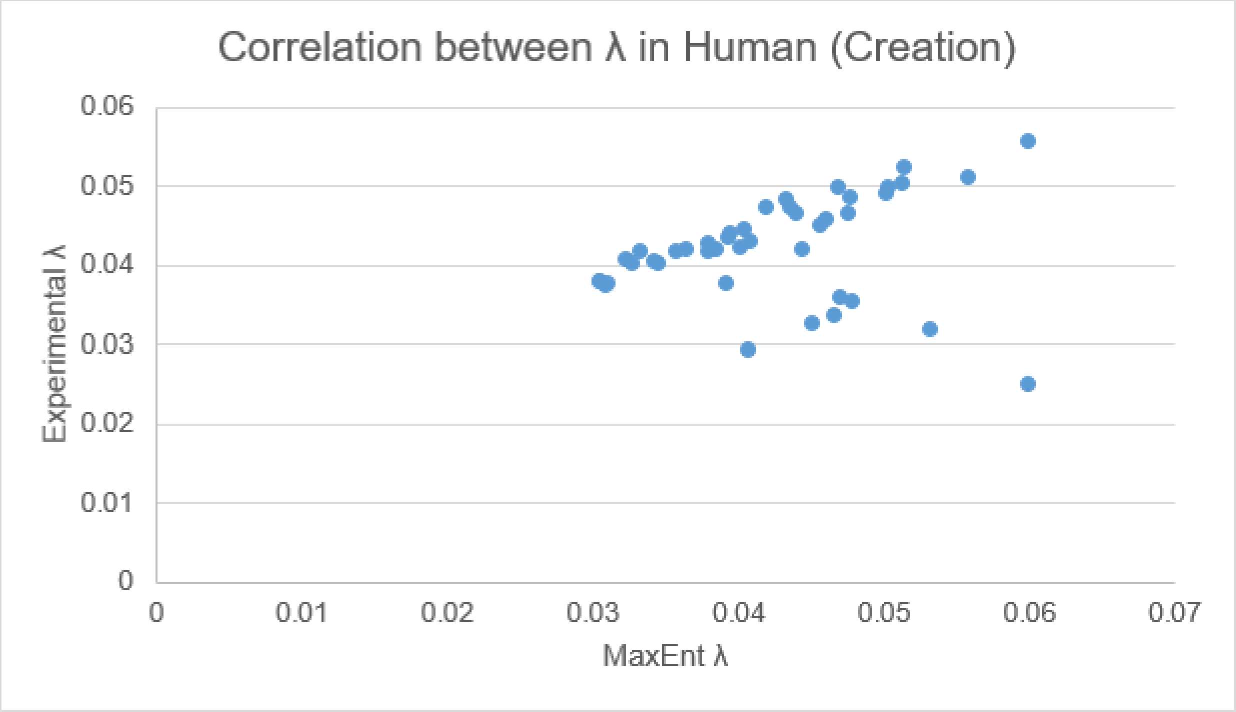
Correlation between λ in Human (Creation)
Experimental λ
0.06
0.05
0.04
0.03
0.02
0.01
0
0.01
0.02
0.03
0.04
0.05
0.06
0.07
MaxEnt λ

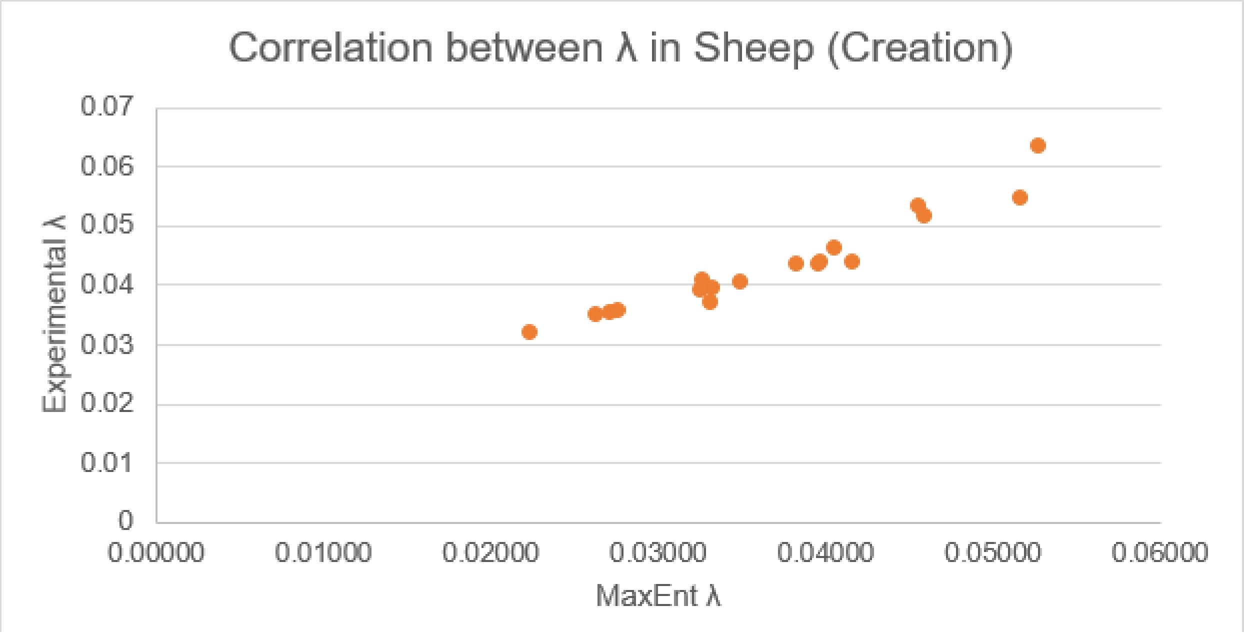
Correlation between λ in Sheep (Creation)
Experimental λ
0.07
0.06
0.05
0.04
0.03
0.02
0.01
0
0.00000
0.01000
0.02000
0.03000
0.04000
0.05000
0.06000
MaxEnt λ

#### S11 – Interpolation

Interpolation was only used in phase visualisation movies, but not in PS detection algorithms. Due to the circular quantity of phase, interpolation was performed using complex phase vectors to avoid incorrect interpolation at phase angle discontinuities [9]. Phase was interpolated from an 8 x 8 grid of electrodes, into a 29 x 29 grid of electrodes wherein each electrode is separated from their neighbour by 3 interpolated points.

#### S12 – MI Surgery for Rat AF

Male Wistar rats weighing 200-275 g were injected subcutaneously (SC) with preoperative buprenorphine (0.03 mg/kg) and anaesthetized with 2% isoflurane. Under endotracheal intubation and assisted ventilation, a left thoracotomy was performed, followed by ligation of the left anterior descending coronary artery with 6-0 silk. The thorax was sutured using a 3-0 silk and the skin was stapled using metal clips. Buprenorphine (0.03 mg/kg) was injected SC 6 and 12 hours postoperatively. Echocardiography was performed at baseline 1 day before the surgery, then 2 weeks later, to assess successful MI, and followed by final echocardiography 3 weeks following surgery. The same day, rats underwent transesophageal electrophysiological study (EPS).
